# Optimizing the power to identify the genetic basis of complex traits with Evolve and Resequence studies

**DOI:** 10.1101/583682

**Authors:** Christos Vlachos, Robert Kofler

**Author notes:** to whom correspondence should be sent.

## Abstract

Evolve and Resequence (E&R) studies are frequently used to dissect the genetic basis of quantitative traits. By subjecting a population to truncating selection for several generations and estimating the allele frequency differences between selected and non-selected populations using Next Generation Sequencing, the loci contributing to the selected trait may be identified. The role of different parameters, such as, the population size or the number of replicate populations have been examined in previous works. However, the influence of the selection regime, i.e. the strength of truncating selection during the experiment, remains little explored. Using whole genome, individual based forward simulations of E&R studies, we found that the power to identify the causative alleles may be maximized by gradually increasing the strength of truncating selection during the experiment. Notably, such an optimal selection regime comes at no or little additional cost in terms of sequencing effort and experimental time. Interestingly, we also found that a selection regime which optimizes the power to identify the causative loci is not necessarily identical to a regime that maximizes the phenotypic response. Finally, our simulations suggest that an E&R study with an optimized selection regime may have a higher power to identify the genetic basis of quantitative traits than a GWAS, highlighting that E&R is a powerful approach for finding the loci underlying complex traits.

## Introduction

Most variation of traits important in agriculture, medicine, ecology and evolution is quantitative (Mackay, 2001). Variation in such quantitative traits (or complex traits) is usually due to multiple segregating loci (Mackay, 2001). For these quantitative traits the simple Mendelian correspondence between genotype and phenotype breaks down, such that one particular phenotype may be due to several distinct genotypes (Lander and Schork, 1994). Unraveling the genetic basis of quantitative traits will be crucial for improving crop yield, leveraging personalized medicine and shedding light on poorly understood evolutionary processes such as extinctions, rapid adaptation and canalization. It has even been argued that identifying the genetic basis of quantitative traits will be the key challenge for biology in the 21*th* century (Mackay, 2001; Losos et al., 2013; Stapley et al., 2010).

Due to this wide interest many approaches for identifying the genetic basis of complex traits have been developed, such as QTL mapping and genome-wide association studies (GWAS) (Mackay, 2001; Korte and Farlow, 2013). These methods suffer from some limitations. QTL studies only capture a limited amount of the variation present in natural populations and the resolution of QTL studies is usually low (Mackay, 2001). Identifying the causative nucleotide (the QTN) is thus rarely achieved with QTL studies (Rockman, 2011). GWAS on the other hand capture more natural variation and have a higher resolution than QTL studies, frequently enabling the identification of some QTNs. GWAS achieve this high resolution by utilizing historical recombination events rather than recombination events occurring within QTL mapping populations (Mackay et al., 2009). However, also GWAS have some limitations. With GWAS it is difficult to identify rare variants and variants of small effect size (Korte and Farlow, 2013; Marchini et al., 2004). Hence alternative approaches for identifying the QTNs are of wide interest.

The advent of Next Generation Sequencing (NGS) made it feasible to monitor adaptation at the genomic level with an approach termed Evolve and Resequence (E&R (Long et al., 2015; Schlötterer et al., 2015)). A base population, that usually captures a substantial amount of the variation of a natural population, is subject to some selective pressure over multiple generations and allele frequency changes are monitored by sequencing the experimental populations. The selective pressure may either be natural, when a population is exposed to a defined environment, or artificial, when a specific phenotype is selected (Schlötterer et al., 2015; Garland and Rose, 2009). Because the selected traits are usually not known in E&R studies relying on natural selection, this approach is mostly used to study the dynamics of adaptation rather than the genetic basis of complex traits. However E&R studies with artificial selection may be a powerful approach for identifying the QTNs, especially since E&R studies rely on both, historical recombination events (base population) and recombination events occurring during the experiment. Using computer simulations several theoretical studies found that E&R studies have a sufficient power to identify selected loci, provided that a powerful experimental design is used (Kofler and Schlötterer, 2014; Baldwin-Brown et al., 2014; Kessner and Novembre, 2015). Such a powerful design usually requires large population sizes (> 1000), several replicates (> 5) and multiple generations of selection (> 90). Hence E&R studies are mostly suitable for small organisms having short generation times such as fruit flies, (Turner et al., 2011; Turner and Miller, 2012; Orozco-Terwengel et al., 2012; Tobler et al., 2014), nematodes (Teotonio et al., 2012), yeast (Kosheleva and Desai, 2017), bacteria (Wannier et al., 2018) and mice (Keightley and Bulfield, 1993). But even for model organisms, E&R studies come at a considerable cost in terms of time and sequencing effort. It is thus important to ensure that the invested resources are optimally utilized by maximizing the power to identify the QTNs. One promising but little explored approach for increasing the performance of E&R studies is the selection regime, i.e. the number of selected individuals during the experiment. This approach is especially promising as an optimized selection regime comes at no, or only little, additional cost (time, sequencing and phenotyping). As a major challenge, an optimal selection regime needs to strike a balance between too weak and too strong selection. With weak selection it will be difficult to distinguish QTNs from neutral loci subject to genetic drift, whereas with strong selection the QTNs may not be distinguished from vast amounts of hitchhikers.

Using genome-wide forward simulations of populations under truncating selection we show that the performance of E&R studies may be maximized by gradually increasing the strength of selection during the experiment, i.e. decreasing the number selected individuals with time. This approach reduces hitchhiking associated with strongly selected loci but nevertheless generates a noticeable response of weakly selected loci. Interestingly, we found that an E&R study with an optimized selection regime may have a higher power to identify QTNs than a GWAS involving several thousands of individuals. A suboptimal selection regime, such as constant strong selection, however results in a poor performance. Our results highlight that the selection regime is a crucial factor determining the success of E&R studies.

## Results

To test if the selection regime has an influence on the performance of E&R studies we performed genome-wide forward simulations with MimicrEE2 (Vlachos and Kofler, 2018). MimicrEE2 is a versatile tool that allows to simulate temporally variable truncating selection with a quantitative trait. We aimed to capture the genomic landscape of *D. melanogaster*, a commonly used model organism for E&R studies (Long et al., 2015; Schlötterer et al., 2015). We used the recombination rate estimates of Comeron et al. (2012) and a base population consisting of 1000 diploid genomes that reproduce the pattern of natural variation found in a *D. melanogaster* population from Vienna (Kofler and Schlötterer, 2014; Bastide et al., 2013) (supplementary fig. 1). Simulations were performed for the major autosomes, where low recombining regions were excluded as they inflate the false positive rate (Kofler and Schlötterer, 2014) (supplementary fig. 1). Based on the recommendations of Kofler and Schlötterer (2014) we simulated an E&R study with a population size of 1000, 10 replicates and 90 generations of selection (table 1). We simulated a quantitative trait model, where 100 randomly selected loci contribute to a trait (100 QTNs). Only loci with frequencies between 5 and 95% where selected, which ensures that all selected SNPs contribute at least moderately to the genotypic variance (Falconer and Mackay, 1960).

**Table 1:** Overview of the default parameters used for the simulations

| parameter | default value |
| --- | --- |
| population size (N) | 1000 |
| number of causative loci | 100 |
| number of generations | 90 |
| replicates | 10 |
| distribution of effect sizes | gamma with shape=0.42 and scale=1 |
| heritability | 1 (genotype to phenotype mapping=1:1) |
| recombination map | Comeron et al. (2012) |
| repetitions | 10 (using different causative loci and effect sizes) |

The effect sizes of the QTNs followed a gamma distribution that captures the distribution of effect sizes found with QTL studies (Hayes and Goddard, 2001; Meuwissen et al., 2001) (table 1). We thus simulated quantitative traits with few large effect loci and many weak effect loci. The sign of the effect (*a* vs. −*a*) was randomly chosen and a heritability of *h*^2^ = 1 was initially used. Simulations for each experimental design were repeated 10 times using different sets of randomly drawn QTNs (random position and effect size) (table 1). The significance of the allele frequency differences between the base population and the evolved populations was estimated with the Cochran-Mantel-Haenszel test (cmh-test)(Landis et al., 1978). This test takes replicates into account and has a good performance with E&R studies (Kofler and Schlötterer, 2014; Schlötterer et al., 2015). The power of the different selection regimes was assessed using Receiver Operating Characteristic (ROC) curves (Hastie et al., 2009), which relate the true-positive (TPR) to the false-positive rates (FPR). The *TPR* can be calculated as *TP/TP* + *FN* where *TP* stands for true positives and *FN* for false negatives. The *FPR* can be calculated as *FP/TN* + *FP*, where *FP* refers to false positives and *TN* to true negatives. A ROC curve having a TPR of 1.0 with a *FPR* of 0.0 indicates the best possible performance. As we are mostly interested in identifying QTNs at a low FPR we used a *FPR* threshold of 0.01 and computed the area under the partial ROC curve 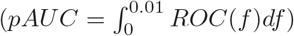 to assess the performance of a selection regime. For an overview of the simulation pipeline see supplementary fig. 2.

### The selection regime influences the performance of E&R studies

We first tested the hypothesis that selection regimes have a significant influence on the performance of E&R studies. We generated different selection regimes by varying the strength of truncating selection throughout the experiment. With truncating selection the individuals having the most pronounced phenotypes are selected and allowed to mate. The offspring of the selected individuals will constitute the next generation. We evaluated the performance of three different selection regimes: a) constant strength of selection, where 50% of the individuals are selected at each generation (50%); b) linearly increasing strength of selection, where 90% of the individuals are selected at the beginning of the experiment and 10% at the end (90 → 10%); and c) linearly decreasing strength of selection, where 10% of the individuals are selected at the beginning of the experiment and 90% at the end (10 → 90%, fig. 1A). Note that the total number of selected individuals is identical for the three selection regimes.

**Figure 1:**
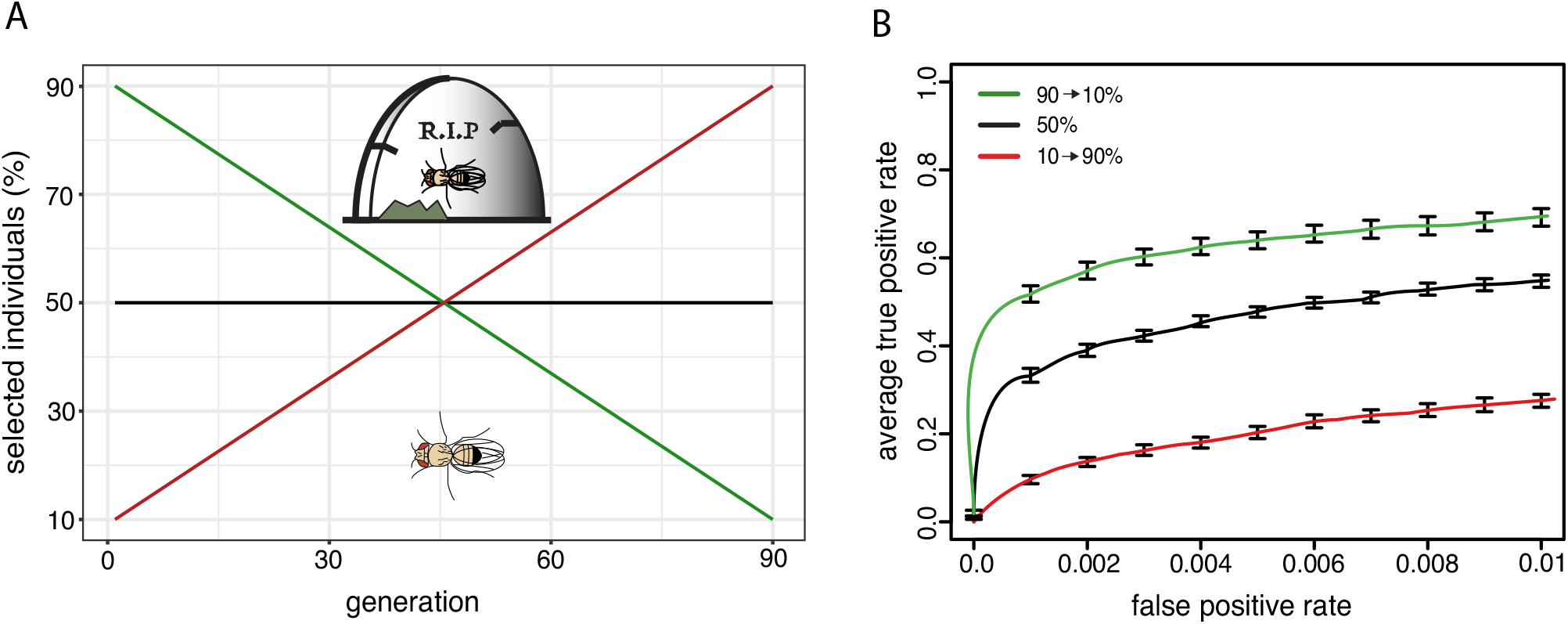
The selection regime has a significant influence on the performance of E&R studies. A) We simulated three truncating selection regimes. The strength of selection increased (green), remained constant (black) or decreased (red) during the experiment. Note that the total number of selected individuals is identical for the three selection regimes. B) ROC curves showing the performance of the three selection regimes. The increasing selection regime has the best performance.

We found that the selection regime has a significant influence on the power to identify the causative loci (Kruskal–Wallis rank-sum test with pAUC; *p* = 2.8e − 06) where linearly increasing strength of selection (henceforth “increasing regime”) had the best performance (fig. 1B). A constant strength of selection had an intermediate performance (henceforth ” constant regime”) and decreasing the strength of selection (henceforth “decreasing regime”) had the worst performance (fig. 1B).

This raises the question why the increasing regime performed better than the constant and the decreasing regime. Apart from technical problems (e.g. sequencing) an E&R study has two sources of noise: genetic drift and hitchhiking of neutral alleles linked to selected loci. If selection is weak, allele frequency changes of neutral loci subject to genetic drift may be more pronounced than the response to selection of the QTNs. On the other hand if selection is strong, alleles linked to selected loci will have little opportunity to recombine to neutral haplotypes. These hitchhikers will thus show a significant response to selection. An optimal selection regime must thus aim to minimize both sources of noise, hitchhiking and drift.

To identify possible causes for the performance differences among selection regimes we investigated the trajectories of strong (effect size > 1), weak (effect size ≤ 1) and neutral alleles in a single replicate of each selection regime (supplementary fig. 3). Most selected loci got fixed (frequency of 1.0, polarized to selected allele) in all three selection regimes, but fixation of selected alleles appears to be delayed in the increasing regime relative to the other regimes (supplementary fig. 3). Delayed fixation of selected alleles could lead to fewer hitchhikers and thus account for the performance differences among selection regimes. To test this hypothesis we quantified the response to selection of strong and weak effect loci in the selection regimes using 50 simulations with different random sets of QTNs (50 sets of QTNs; 1 replicate; supplementary fig. 4). Fixation of both strong and weak effect loci was significantly delayed in the increasing regime compared to the constant and the decreasing regime (Wilcoxon rank-sum test; *p* < 2.2e − 16). Furthermore, fixation of strong effect loci was more delayed than fixation of weak effect loci (strong: *incr./cons*. = 1.9*x, incr./decr*. = 5.2*x*; weak: *incr./cons*. = 1.4*x, incr./decr*. = 3.3*x*; supplementary fig.4). This suggests that the increasing regime affords more time to neutral alleles to recombine away from selected haplotypes.

Next, we aimed to quantify the extend of hitchhiking for the three selection regimes. Replicated E&R studies allow to roughly distinguish between hitchhikers and alleles subject to genetic drift by the consistency of the allele frequency change among replicates. For example, alleles that increase in frequency in some replicates but decrease in frequency in others are likely subject to drift, whereas alleles that increase in frequency in all replicates are likely hitchhikers. We thus classified loci as hitchhikers when the allele frequency consistently changed in the same direction in all 10 replicates (excluding QTNs). In agreement with our hypothesis we found fewer hitchhikers in the increasing regime than in the other two regimes (10 set of QTNs; 10 replicates; supplementary fig. 5). Furthermore, the allele frequency change of hitchhikers was least pronounced in the increasing regime (supplementary fig. 5). Of course, constant weak selection where for example 90% of the individuals are selected may reduce hitchhiking even more than the tested increasing regime (supplementary fig. 4). However constant weak selection results in a overall reduced response to selection (supplementary fig. 4) such that the noise generated by genetic drift may dominate the weak signal of selected loci. In summary, we propose that the increasing regime has a high performance because it initially delays the fixation of strong effect loci but amplifies the response of weak effect loci at the end of the experiment.

So far we evaluated the performance of three different selection regimes having an identical total number of selected individuals. However many more different selection regimes, with varying amounts of selected individuals are feasible. We thus carried out additional simulations to test if increasing regimes outperform other selection regimes.

### Linearly increasing the strength of selection maximizes the power to identify QTNs

Among all feasible linear selection regimes (increasing, constant and decreasing) we aimed to identify the regime that results in the highest power to identify the QTNs. Since genome-wide forward simulations are computationally demanding we needed to limit the number of necessary simulations. As the decreasing regime had a poor performance we solely considered increasing and constant regimes (fig. 1; supplementary fig. 4,5). Furthermore we evaluated the performance of selection regimes in steps of 10% selected individuals (fig. 2A,B). To identify the best increasing regime we thus evaluated the performance of 36 different regimes (90 → 80%, 90 → 70%,…,90 → 10%,…,20 → 10%; 2 A, left panel). Based on the area under the ROC curve (*pAUC*) we found that the increasing regime where 90% of the individuals are selected at the beginning of the experiment and 20% at the end had the best performance (90 → 20% increasing regime; fig. 2A). Out of the constant regimes, on the other hand, selection of 80% of the individuals resulted in the highest power to identify the QTNs. (fig 2B). This is in agreement with the results of Kessner and Novembre (2015) who evaluated the performance of multiple constant regimes (20%, 40%, 60% and 80%) under a QTL model with truncating selection and found that selection of 80% of individuals performed best. However, we found that the best increasing regime had a higher power to identify QTNs than any of the evaluated constant regimes (fig. 2B; Wilcoxon rank-sum test with pAUC; 90 → 20% vs each constant regime; *p* ≤ 0.0002).

**Figure 2:**
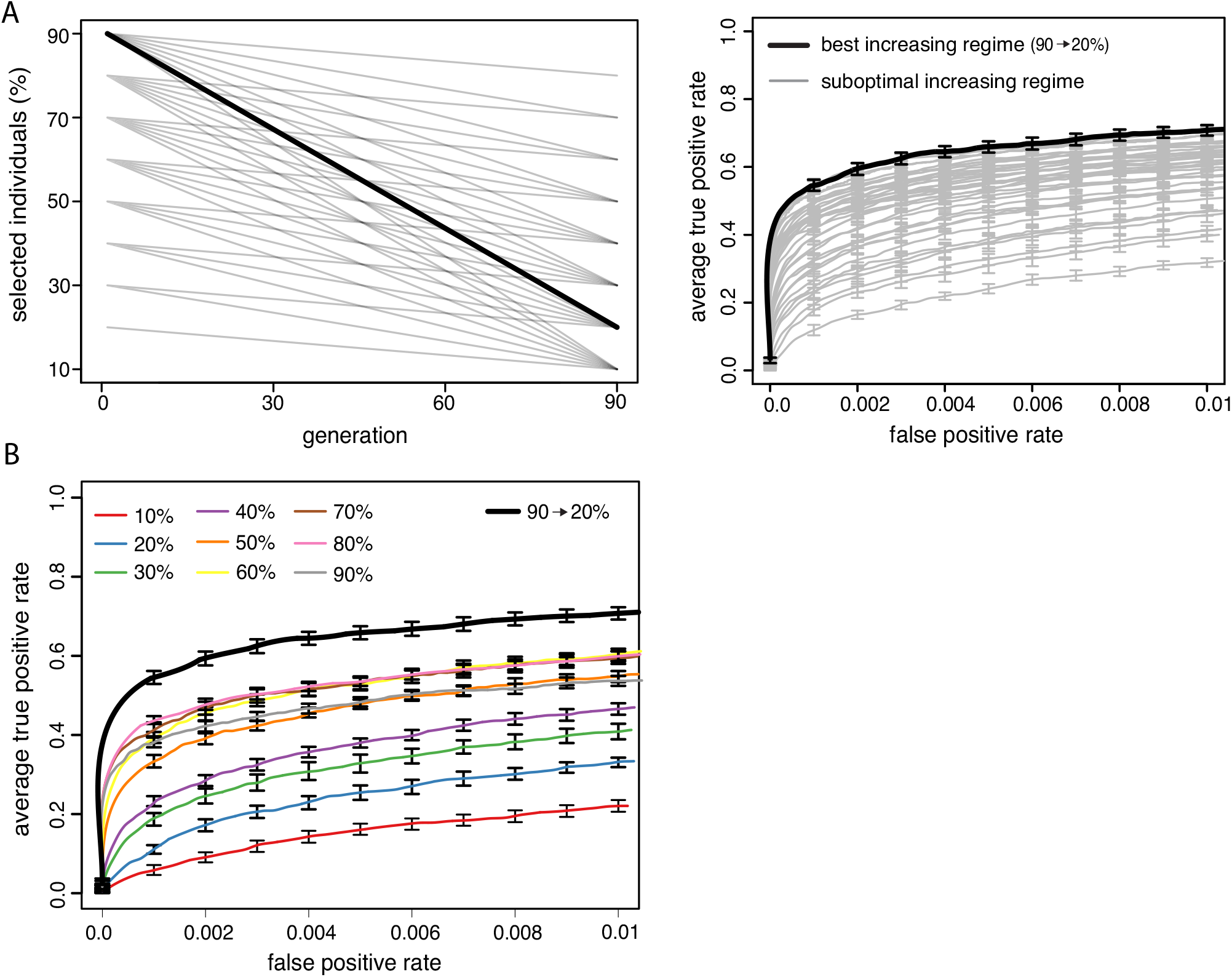
Increasing the strength of selection during an E&R study enhances the power to identify QTNs. A) Approach for identifying the best increasing regime. We evaluated the performance of all possible increasing regimes using steps of 10% (left panel). The best selection regime (bold) has the largest area under the ROC curve (right panel). B) The best increasing regime outperforms the evaluated constant regimes (steps of 10%)

### Influence of the experimental design

Depending on the experimental organism, the default of 90 generations of selection may be quite time consuming (e.g. 3 years in Drosophila). We thus asked whether a high performance may also be achieved with shorter experiments if an optimized selection regime is used. We evaluated the performance of different selection regimes for 20, 45 and 90 generations of selection (fig. 3). In agreement with previous works we found that performance of E&R studies increases with the number of generations of selection (Kofler and Schlötterer, 2014; Kessner and Novembre, 2015; Baldwin-Brown et al., 2014) (fig. 3). Of the constant regimes, selection of 80% of the individuals consistently had the best performance, irrespective of the length of the experiment (fig. 3). This is again in agreement with Kessner and Novembre (2015). With 90 and 45 generations of selection the best increasing regime significantly outperformed the constant regimes (Wilcoxon rank sum test with pAUC; *p*_45_ = 0.014, *p*_90_ = 0.0002). With 20 generations of selection, however, the performance of the best increasing and constant regime was quite similar (Wilcoxon rank sum test with pAUC; *p*_20_ = 0.68). We also noticed that the optimal increasing regime changed from 90 → 20% with 90 generations of selection to 90 → 70% with 20 generations. With 45 generations of selection the performance of the 80 → 10% increasing regime was not significantly different from the 90 → 20% increasing regime (Wilcoxon rank sum test, *p* = 1). For short experiments the performance of the best increasing regime thus approaches the performance of the best constant regime. Short experiments may not provide sufficient time for the benefit of increasing regimes, such as the delayed fixation of large effect loci, to take effect and thus influence the performance. We conclude that an optimized selection regime is not able to compensate for the loss of performance incurred by reducing the generations of selection. In fact the advantage of an optimized selection regime, such as the increasing regime, is most pronounced for long experiments.

**Figure 3:**
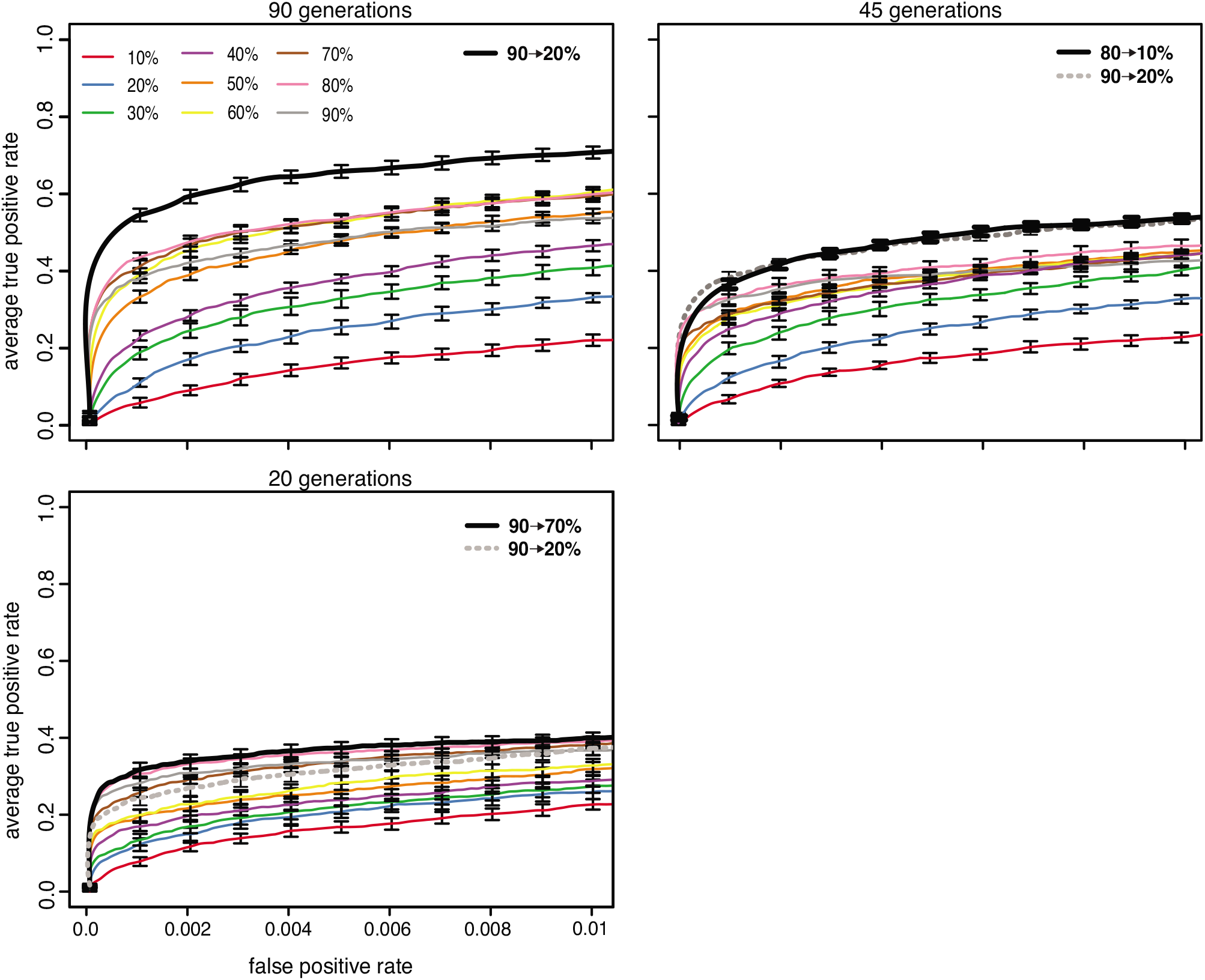
Influence of the number of generations (*default* = 90). The performance of the best increasing regime (black, bold), the 90 → 20% increasing regime (grey, dashed) and the constant regimes (different colors) is shown. Although increasing regimes consistently perform best, the advantage of an increasing regime is most pronounced for long E&R studies.

Next, we assessed the effect of the number of replicate populations on the shape of the optimal selection regime. We evaluated the performance of the different selection regimes with 3, 5 and 10 replicates (10 is the default). Our results suggest that the best increasing regime significantly outperforms the best constant regime when 10 or 5 replicates are used (Wilcoxon rank sum test with pAUC; *p*_5_ = 0.0004, *p*_10_ = 0.0002; supplementary fig. 6). For 3 replicates the performance of the best increasing regime is not significantly different from the best constant regime (Wilcoxon rank sum test with pAUC; *p*_3_ = 0.5). The advantage of an increasing regime is thus most pronounced when many replicates are used. Interestingly, also the shape of the optimal selection regime depends on the number of replicates, where strong selection at the of end the experiment is especially beneficial when many replicates are used (supplementary fig. 6).

This raises the question why replication influences the shape of the optimal selection regime. For loci mostly subject to genetic drift the direction of the allele frequency change will vary across replicates. As high cmh-scores require consistent allele frequency changes across replicates, it will be easier to distinguish between selected loci and loci subject to drift when many replicates are used. An elevated strength of selection at the end of highly replicated E&R studies may thus boost the response to selection of weak effect loci without incurring excessive additional noise from genetic drift caused by the population size reduction at the end of the experiment.

Finally, we investigated the influence of the population size (supplementary fig. 7). We found that an increasing regime consistently performed better than the constant regimes where especially the 90 → 20% increasing regime had a high performance with the evaluated population sizes (supplementary fig. 7).

To summarize, in all tested experimental designs an increasing regime outperformed the constant regimes. However, the slope and intercept of the optimal increasing regime depends on the experimental design where especially the number of replicates and the length of the experiment had a noticeable influence.

### Influence of the trait architecture

We next asked if the optimal selection regime depends on the architecture of a quantitative trait. We investigated the influence of the number of QTNs, the heritability and the effect size distribution of the QTNs.

First, we considered the influence of the number of QTNs. We simulated E&R studies with 25 and 1000 QTNs in addition to the default of 100 QTNs. In agreement with previous works we found that the performance of E&R studies is weak when the number of QTNs is large [supplementary fig. 8; (Kofler and Schlötterer, 2014; Kessner and Novembre, 2015)]. A large number of QTNs results in widespread interference among selected loci, which can not (or only very slowly) be resolved by recombination events arising during the experiment. Irrespective of the number of QTNs, of the constant regimes selection of 80% the individuals consistently performed best. This is again in agreement with previous works (Kessner and Novembre, 2015). Although an increasing regime consistently performed better than the constant regimes (supplementary fig. 8) the advantage of an increasing regime was most pronounced for intermediate numbers of QTNs (100; supplementary fig. 8). We noticed that the 90 → 20% increasing regime resulted in a good performance for diverse numbers of QTNs (supplementary fig. 8).

The heritability, i.e. the proportion of the phenotypic variance that is due to the genotype, varies among environments, populations and traits (Visscher et al., 2008; Falconer, 1992). To explore the influence of the heritability we simulated E&R studies with heritabilities of *h*^2^ =0.3 and *h*^2^ = 0.6 in addition to the default of *h*^2^ = 1.0 (supplementary fig. 9). We found that an increasing regime consistently outperformed the constant regimes (supplementary fig. 9) (Wilcoxon rank sum test; *p*_0.3_ = 0.035, *p*_0.6_ = 0.035, *p*_1_ = 0.0002). Especially the 90 → 20% increasing regime had a high performance across different heritabilities. The advantage of an increasing regime was however most pronounced for a high heritability (*h*^2^ = 1; supplementary fig. 9). We also noticed that the influence of the selection regime diminishes with decreasing heritability (supplementary fig. 9).

Finally we evaluated the influence of the effect size distribution of the QTNs. Per default we used a distribution that captures the effect sizes found in QTL studies (gamma distribution with shape 0.42) (Hayes and Goddard, 2001; Meuwissen et al., 2001). However, the effect size distribution may vary among traits, populations and even environments (Dittmar et al., 2016; El-Soda et al., 2014). To evaluate the influence of the effect size distribution we simulated E&R studies with QTNs drawn from different gamma distributions with shape parameters ranging from 0.1 to 1.0. Furthermore we simulated one distribution where all loci had identical effect sizes. Note that the absolute value of the effect size is not important when truncating selection (i.e. soft selection) is used and that effect sizes are getting more similar with an increasing shape parameter (e.g. ratio between the 10% largest and smallest effect sizes Γ_0.1_ = 602, 622; Γ_1.0_ = 62). Interestingly, we found that an increasing regime consistently performed best when effect sizes followed a gamma distribution (Wilcoxon rank sum test with pAUC; *p*_0.1_ = 4.33*e* − 05, *p*_0.42_ = 2*e* − 04, *p*_0.7_ = 3.2*e* − 04, *p*_1.0_ = 0.035; fig. 4). However, when effect sizes were identical constant selection of 90% of the individuals performed best. For gamma distributed effect sizes the 90 → 20% increasing regime consistently had a high performance. Generally we note that increasing regimes perform best when effect sizes are highly unequally distributed (e.g. gamma with shape=0.1). This is in agreement with our proposed explanation for the good performance of increasing regimes, i.e. an initially delayed fixation of large effect loci combined with an encouraged fixation of small effect loci at later generations. Such dynamics will not be beneficial when all loci have identical effect sizes.

**Figure 4:**
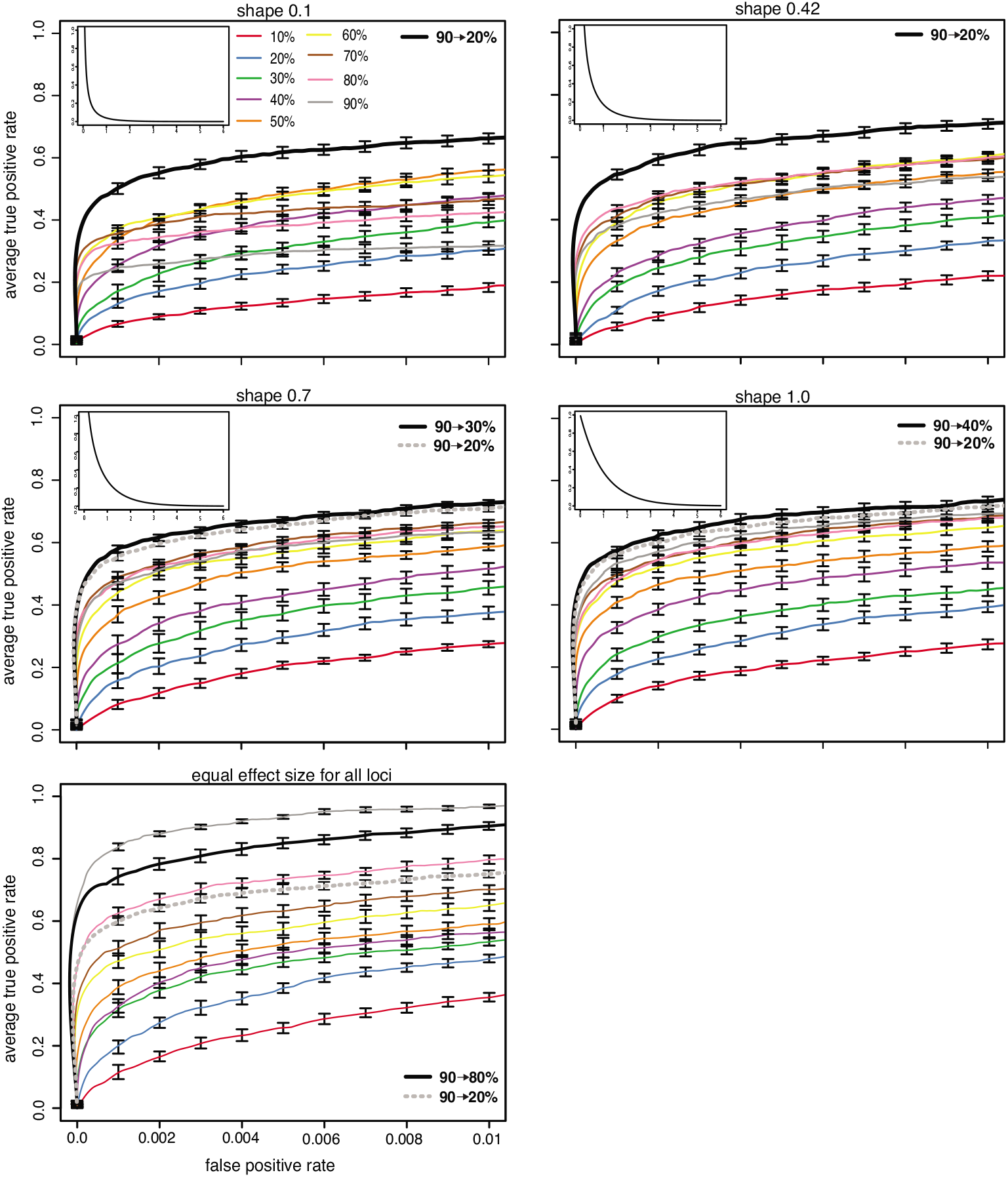
Influence of the effect size distribution. We evaluated the performance of selection regimes with different effect size distributions. In addition to different gamma distributions we included a scenario where all effect sizes were identical (default is gamma with shape=0.42). Note that effect sizes become more similar with an increasing shape parameter and that the advantage of the increasing regime is most pronounced for highly unequal effect sizes (e.g. *shape* = 0.1).

Finally, we asked if selection regimes may influence our ability to estimate the effect size distribution of QTNs. We investigated the effect sizes of QTNs among the 2000 most significant SNPs in the simulations with gamma distributed effect sizes. (supplementary fig. 10) For both, increasing and constant regimes, the 2000 most significant SNPs only contained a fraction of the QTNs (supplementary fig. 10). However the best increasing regime allowed to recover a higher fraction of the QTNs than the best constant regime (increasing regimes 42.3%, constant regimes 29.2%). Especially loci with weak effect sizes were more readily identified with an increasing regime (supplementary fig. 10). Hence, increasing regimes may enable us to more accurately recover the effect size distribution of QTNs than constant regimes.

To summarize, with the exception of a trait architecture where QTNs have identical effects, increasing regimes outperformed constant regimes over a wide range of different trait architectures.

### Selection that optimizes the power to identify QTNs does not necessarily maximize the phenotypic response

Before the advent of E&R studies truncating selection was used to change a phenotype of interest. In a classic example the oil content of maize was raised from 5% to about 20% by continuously selecting the individuals with the highest oil content (Dudley and Lambert, 2010). We were interested whether selection regimes that aim to optimize the power to identify QTNs (henceforth “QTN regime”) are identical to selection regimes that aim to maximize the phenotypic response (henceforth “phenotype regime”). To address this question we simulated multiple truncating selection regimes in steps of 10% selected individuals and identified i) the best QTN regime (see above) and ii) the best phenotype regime, i.e. the regime that maximizes the phenotypic difference between the base and the evolved population (*R*).

We found that a selection regime that maximizes the phenotypic response to selection does not necessarily have a high power to identify QTNs (fig. 5). This discrepancy is especially pronounced for short experiments (fig. 5; 20 generations; Wilcoxon rank sum test; 90 → 70% vs 20 → 10%; *p* = 2.2*e*−16). In order to maximize the phenotypic response strong selection is optimal for short experiments whereas weaker selection is best for long experiments (fig. 5). This is in agreement with previous theoretical works which found that the optimal strength of selection decreases with the length of the experiment (Robertson, 1970a). For very long experiments (infinite generations) and in the absence of linkage the optimal phenotype regime approaches constant selection of 50% of the individuals (Robertson, 1960). With linkage, as in our simulations, a slightly larger fraction of selected individuals is optimal (Robertson, 1970b). Previous works also found that for finite experiments an increasing regime (albeit a sigmoid increase in the strength of selection) yields the largest phenotypic response to selection (Robertson, 1970a). Interestingly with QTN regimes the situation is reversed. Here, weak selection is optimal for short experiments whereas stronger selection, especially at the end of the experiment, is best for long E&R studies (fig. 5). Due to these two contrasting trends the optimal QTN and phenotype regime are very similar at intermediate generations of selection (45 generations; fig. 5A).

**Figure 5:**
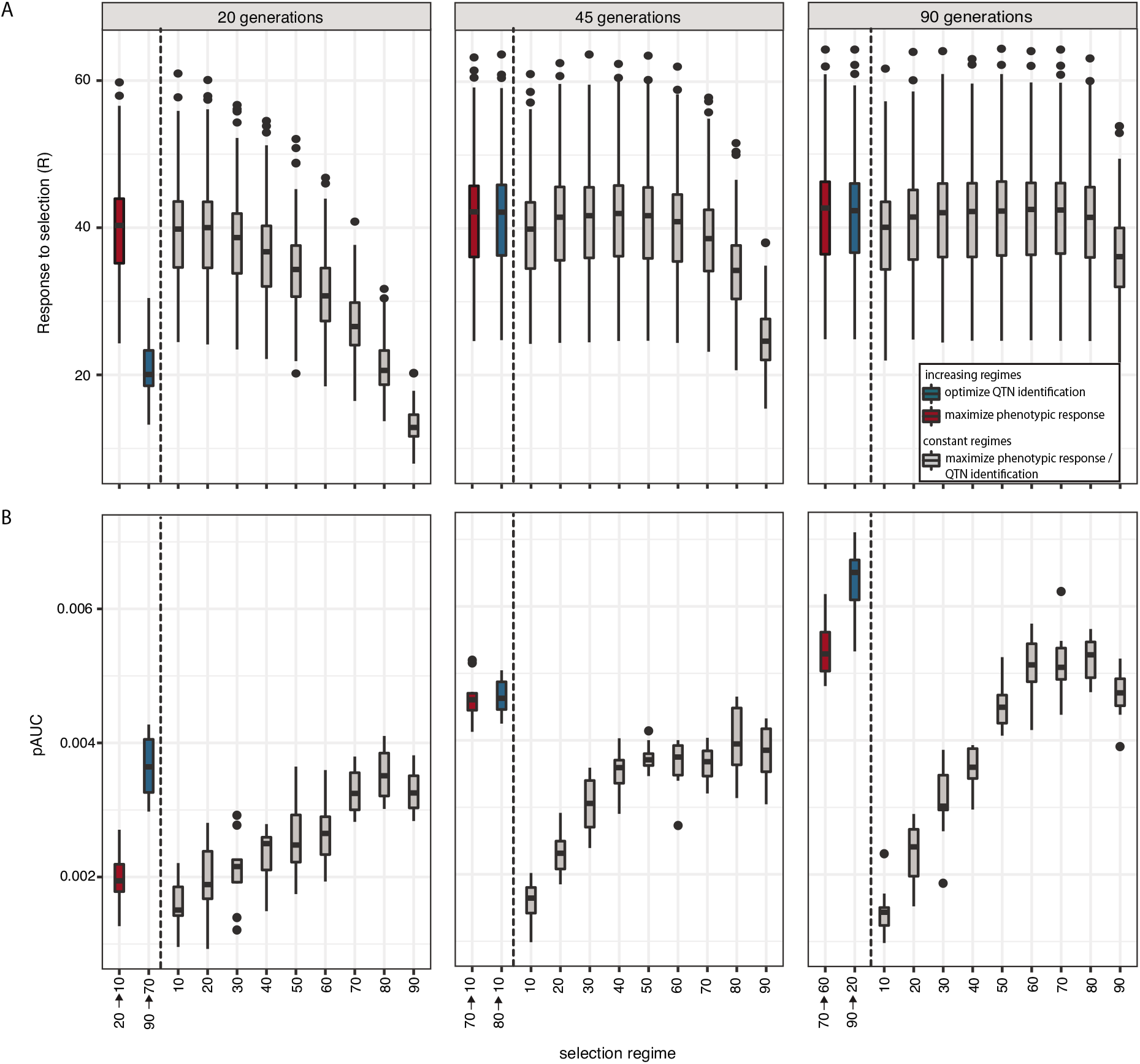
Selection regimes that maximize the power to identify QTNs (red) are not necessarily identical to regimes that maximize the phenotypic response to selection (blue). As a reference the performance of constant selection regimes (grey) is shown. A) Phenotypic response to selection. B) Power to identify the QTNs assessed by the partial area under the ROC curve (highest possible performance *pAUC* = 0.01).

What is responsible for the discrepancy between optimal QTN and phenotype regimes? We think that two factors are likely important: hitchhiking and replication. Neutral hitchhikers are a major source of noise for QTN studies but do not impede selection of phenotypes. Hence, for short studies, phenotype regimes may benefit more from strong selection than QTN regimes. On the other hand, the phenotypic response to selection is not affected by the number of replicates but the power to identify QTNs increases with replication (supplementary fig. 6). With QTN studies, strong selection at the end seems to be especially beneficial when many replicates are used (supplementary fig. 6). This influence of replication may explain why strong selection at the end is more beneficial for QTN regimes than for phenotype regimes.

We however also made the observation that an optimized selection regime seems to be more important for QTN identification than for phenotype selection. Even suboptimal phenotype regimes yield a substantial phenotypic response to selection (fig. 5A; most linear regimes yield a similar *R*). This is in agreement with previous works which found that the phenotypic response to selection is quite robust over many different selection regimes (Robertson, 1960). However the same observation does not hold for QTN regimes, where deviations from the best regime lead to a noticeable drop in the power to identify QTNs (fig. 5B; linear regimes lead to dissimilar *pAUC* values).

We conclude that selection regimes that maximize the phenotypic response to selection are not necessarily identical to regimes that have a high power to identify QTNs. Furthermore, optimizing the selection regime is more important for QTN identification than for phenotype selection.

### E&R versus GWAS

One of the most widely used approaches for identifying the genetic basis of quantitative traits are genome-wide association studies (GWAS) (Visscher et al., 2012). They have for example been used to shed light on the genetic basis of schizophrenia in humans (Ripke et al., 2014) and starvation resistance in Drosophila (Mackay et al., 2012). GWAS allow to more accurately pinpoint the location of the causative variants than the widely used QTL studies (Mackay, 2001). GWAS achieve this high resolution by utilizing historical recombination events while QTL studies solely rely on recombination events occurring in the mapping populations (Mackay et al., 2009). Since E&R studies utilize both historical recombination events and recombination events occurring in the experimental populations we hypothesized that E&R may offer a higher power to identify QTNs than GWAS.

Ideally, one would compare the performance of a GWAS to an E&R study requiring identical effort, in terms of phenotyping, time and sequencing. Such a comparison is however difficult to accomplish. While GWAS typically require sequencing and phenotyping of each strain/genotype separately, shortcuts may be used for E&R studies. For example, most E&R studies solely require estimates of allele frequencies which can be readily obtained by sequencing the populations as pools (e.g. with 10 replicates only 20 sequencing libraries are necessary). With E&R studies shortcuts may also be used for phenotyping. For example, Turner et al. (2011) performed an E&R study selecting for increased body size in Drosophila using a sieving apparatus.

We therefore decided to compare the performance of an E&R study (using default parameters; table 1) to multiple GWAS having different population sizes (ranging from 500 to 8000). By iteratively sampling haplotypes of a small population from the next larger population we ensured that SNPs segregating in small populations are a subset of SNPs segregating in each of the larger populations. Furthermore, solely SNPs with a frequency between 5% and 95% in each population were picked as QTNs (supplementary fig. 11). GWAS was performed with the widely used tool SNPtest (Marchini et al., 2007) and the performance of the two approaches was evaluated with ROC curves (fig. 6). Note that ROC curves avoid the problem of picking arbitrary significance thresholds for GWAS and E&R.

**Figure 6:**
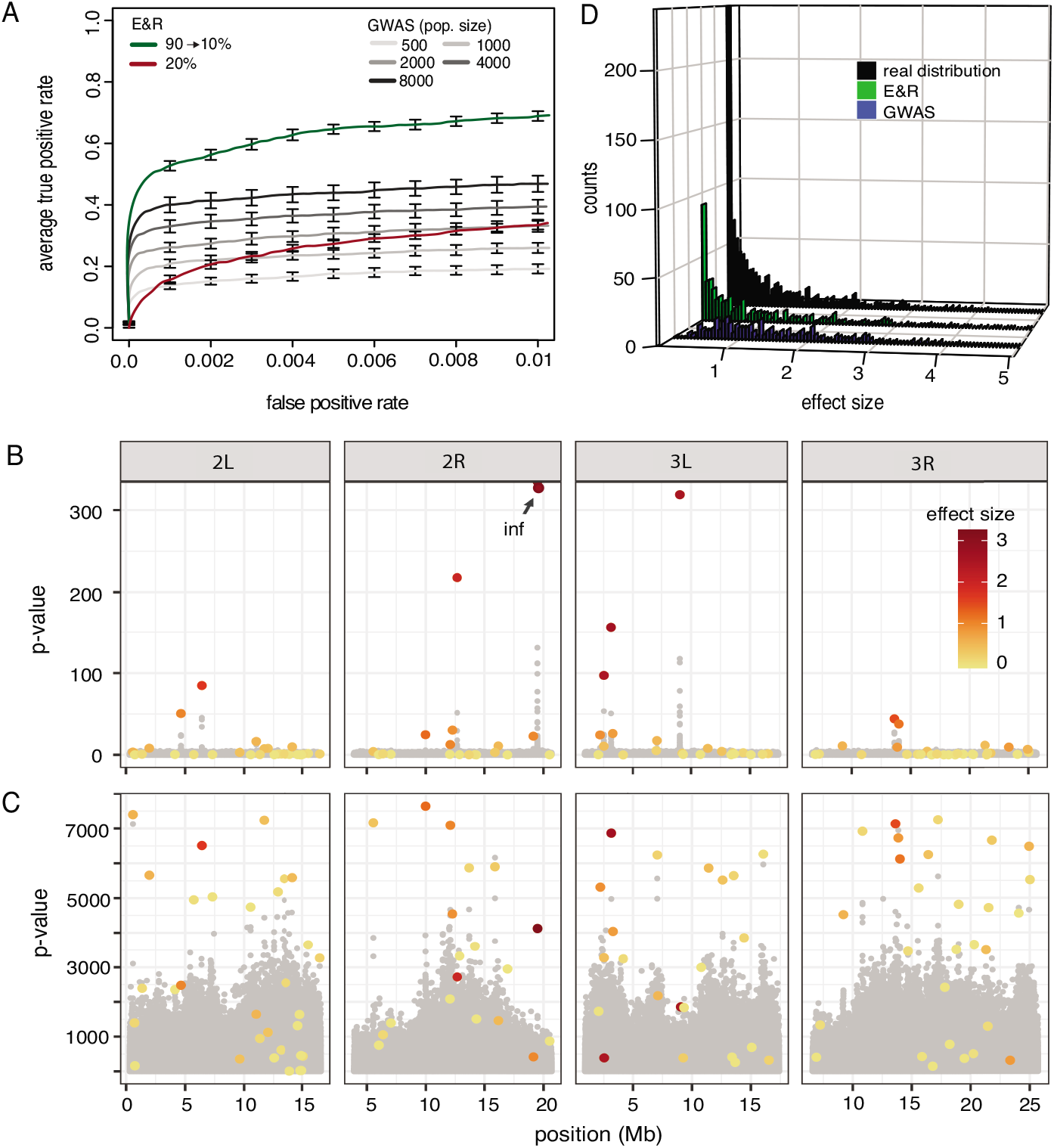
Performance of E&R and GWAS. A) Performance of GWAS with different population sizes and E&R studies using a powerful (green; 90 → 10%) and a suboptimal selection regime (red; constant 20%). The E&R study with an optimized selection regime has a higher power to identify QTNs than the evaluated GWAS. B) Manhattan plot for a GWAS (*N* = 8000). The effect sizes of QTNs are shown in a color gradient, where large effect loci are red. C) Manhattan plot for a E&R study (*N* = 1000, powerful selection regime) D) Histogram of the effect sizes recovered with GWAS and E&R. The 2000 most significant SNPs were used for each approach. The GWAS recovered mostly loci of large effect size while the E&R also recovered many loci of small effect. A summary over 10 experiments is shown.

As expected, the power of GWAS increased with the population size (fig. 6A) (Gibson, 2018). Interestingly, an E&R study with an optimized regime (90 → 10%) had a higher power to identify QTNs than a GWAS with 8000 individuals (fig. 6A); (Wilcoxon rank sum test; E&R_1000_ vs GWAS_8000_; *p* = 2.16*e* − 05). This is also evident from the Manhattan plots, where most peaks in the GWAS are due to large effect loci while also many small effect loci generate peaks in the E&R study (fig. 6B,C). However, the power of E&R drops dramatically when a suboptimal selection regime is used (fig. 6C; red, 20% constant selection), highlighting that the selection regime is a crucial factor determining the performance of an E&R study.

Next we tested if the performance differences between E&R studies and GWAS depend on the architecture of a trait. With a reduced heritability (*h*^2^ = 0.5) the E&R study also had a higher performance than the GWAS (Wilcoxon rank sum test; E&R_1000_ vs GWAS_8000_; *p* = 0.023; supplementary fig. 12). Interestingly, when all loci have identical effect sizes, the GWAS had a higher performance than the E&R study (Wilcoxon rank sum test; E&R_1000_ vs GWAS_8000_; *p* = 1.08*e* − 05; supplementary fig. 12). One complication however arises from the fact that large populations have more polymorphism than small populations. Thus for a given FPR threshold different numbers of false positive SNPs will be compared. We therefore repeated this analysis using absolute numbers of false positive SNPs, but obtained largely similar results (supplementary fig 13).

It has been shown that GWAS have a low power with rare alleles and alleles of small effect (Visscher et al., 2012; Gibson, 2011). We were thus interested if E&R studies suffer from the same or similar weaknesses and investigated the effect sizes and allele frequencies of QTNs among the 2000 most significant SNPs identified with either approach.

Consistent with expectations, the GWAS (*N* = 8000) identified all large effect loci (> 1: 100%) but only few of the small effect loci (≤ 1: 30%; fig. 6D). An E&R study with an optimized design had a worse performance with large effect loci (> 1: 64%) but a higher performance with small effect loci (≤ 1: 50%; fig. 6D). Since small effect loci were more abundant than large effect loci, the E&R study allowed to identify significantly more QTNs than the GWAS (Wilcoxon rank sum test; E&R_1000_ vs GWAS_8000_; *p* = 0.0008).

Furthermore, the GWAS identified most of the loci with intermediate allele frequencies (0.2 ≥ *f* ≥ 0.8: 46%) but few of the alleles with low or high frequencies (*f* < 0.2: 32%, *f* > 0.8: 36%; supplementary fig. 14). By contrast, the E&R study allowed to identify most of the loci with low and medium frequencies in the base population (*f* < 0.2: 90%, 0.2 ≥ *f* ≥ 0.8: 57%) but few of the loci having a high frequency (*f* > 0.8: 0%; supplementary fig. 14). This is due to the fact that selected loci already starting at a high frequency may only exhibit a small allele frequency change, which will usually result in insignificant p-values (low signal). So far, we simulated an E&R study with a powerful design (*N* = 1000, 10 replicate, 90 generations). Although feasible with organisms such as Drosophila (e.g. Graves et al. (2017); Barghi et al. (2019)), this design requires a substantial effort and may therefore be out of reach for many research questions. We were thus interested in the performance of E&R studies with a less powerful design (*N* = 500, 5 replicates, 45 generations). This low-budget design still resulted in a considerable power to identify QTNs, with a performance comparable to a GWAS with about 1000 to 2000 individuals (supplementary fig. 15).

We conclude that an optimized E&R study may provide a higher power to identify QTNs than a GWAS. E&R studies avoid some problems of GWAS, such as a low power with rare alleles and alleles of weak effect, but have other weaknesses in turn, such as a low power with alleles starting at high frequency and a sensitivity to suboptimal selection regimes.

## Discussion

We showed that E&R is a powerful approach for dissecting the genetic basis of complex traits and that the performance of E&R can be optimized by gradually increasing the strength of selection during the experiment. In contrast to previous works which showed that the performance of E&R studies may be improved by increasing the number of replicates, the length of the experiment and the population size (Baldwin-Brown et al., 2014; Kofler and Schlötterer, 2014; Kessner and Novembre, 2015), an optimized selection regime as suggested in this work comes at no, or only little, additional cost.

All approaches for identifying the genetic basis of complex traits rely on a crucial assumption about the distribution of effect sizes. A classic model proposed by Fisher and others holds that an infinite number of loci with equal and small effects contribute to a quantitative trait (Fisher, 1930; Barton et al., 2017). If this model is correct any attempts to identify the QTNs, irrespective of the used method (e.g. GWAS or E&R), are hopeless (Mackay, 2001). Alternatively, Robertson (1970a) and others suggested that the distribution of QTN effects may be resembling an exponential distribution, with few loci having large effects and many loci having small effects (Mackay, 2001). In this case it should be feasible to identify at least a fraction of the QTNs (Mackay, 2001). While there is evidence that the infinitesimal model is a good approximation for many traits, there is also substantial evidence that effect sizes of many traits follow a more or less exponential distribution, with a few QTNs of large and many QTNs of small effect. (Mackay, 2001; Hayes and Goddard, 2001). For these traits it should be feasible to identify QTNs, at least QTNs with an appreciable effect size. We thus assumed a finite architecture of quantitative traits (10 to 1000 QTNs; mostly using gamma distributed effect sizes) throughout the manuscript. The assumption of a limited number of QTNs also explains why we did not simulate *de novo* mutations. Under an infinitesimal model most mutations will hit a QTN and thus affect the quantitative trait. Hence, *de novo* mutations will generate some genetic variation at each generation (Barton et al., 2017). Under the alternative assumption of a finite trait architecture *de novo* mutations will rarely hit one of the few QTNs and thus only generate a limited amount of genetic variation. Furthermore, even if *de novo* mutations hit a QTN, the mutation will be restricted to a single replicate and thus only have a minor influence on the dynamics of highly replicated E&R studies as simulated in this work. For these reasons we did not consider *de novo* mutations.

Here, we aimed to identify the selection regime that maximizes the power to identify QTNs. Since genome-wide forward simulations are computationally demanding we simulated increasing regimes using steps of 10% selected individuals. Due to this discrete sampling of selection regimes we likely missed the absolutely best increasing regime. Nevertheless we think that our approach allowed us to obtain a reasonable approximation of the absolutely best regime as, for example, the three regimes with the highest performance in our simulations consistently have a very similar performance, slope and intercept (supplementary table 1). It is however feasible that non-linear selection regimes achieve a better performance than linear regimes. For example Robertson (1970b) found that the best selection regime for maximizing the phenotypic response to selection has a sigmoid shape. Mostly for computational reasons we did not consider non-linear regimes. Evaluating the different linear regimes already required about 38,000 CPU hours and specifying the shape of non-linear regimes will at least require one additional parameter which would substantially increase the number of necessary simulations.

We assessed the significance of the response to selection using the cmh-test which contrasts, for each SNP, the allele frequency of the base and the evolved populations (Landis et al., 1978). The cmh-test is fast, implemented in user-friendly software and, so far, has the best performance among tests that rely on allele frequency estimates for two time-points (Kofler and Schlötterer, 2014; Schlötterer et al., 2015; Kofler et al., 2011). For theses reasons the cmh-test is widely used in E&R studies (Orozco-Terwengel et al., 2012; Martins et al., 2014; Tobler et al., 2014; Barghi et al., 2019; Phillips et al., 2018; Kelly and Hughes, 2018). Recently however several test-statistics became available that utilize time series data, i.e. allele frequencies estimates for multiple time points (> 2) during the experiment (Topa et al., 2015; Iranmehr et al., 2017; Spitzer et al., 2019). We were interested if our conclusion, that an increasing regime enhances the power to identify QTNs, also holds when a time-series based test-statistic is used. We evaluated an adaptation of the cmh-test to E&R studies, which utilizes time-series data and takes the over-dispersion resulting from drift and pooled sequencing into account (Spitzer et al., 2019). Additionally, we evaluated CLEAR, a composite likelihood based approach for detecting selected regions with E&R studies. With both test statistics the selection regime had a significant influence on the power to identify the QTNs and the increasing regime outperformed the constant regime (supplementary fig. 16, 17). Throughout this work we assumed that allele frequencies were accurately estimated, which usually requires sequencing all individuals in a population separately. As this approach is prohibitively costly most E&R studies rely on Pool-Seq, i.e. sequencing populations as pools, to obtain allele frequency estimates (Schlötterer et al., 2014; Long et al., 2015). To evaluate the influence of the coverage, a crucial parameter determining the accuracy of allele frequency estimates with Pool-Seq, we performed binomial sampling of allele frequencies to different coverages. An increasing regime outperformed the constant regime, irrespective of the coverage (supplementary fig. 18).

We found that the optimal selection regime depends on the experimental setup and the trait architecture. Unfortunately the trait architecture is usually not known at the onset of an E&R study. In fact shedding light on the architecture of a trait of interest may be the aim of an E&R study. We however noticed that the 90 → 20 increasing regime, where 90% of the individuals are selected at the beginning and 20% at the end of the experiment, shows a good performance over a wide range of parameters. Only exceptions are short experiments and traits with QTNs of identical effect sizes, where constant weak selection (e.g. 80% selected individuals) achieves the best results.

A scan of previously used selection regimes showed that many E&R studies applied very strong selection: Turner et al. (2011) selected the ≈ 9% largest flies for over 100 generations; Turner and Miller (2012) selected 20% of the flies with the longest pause in courtship song for 14 generations; Hardy et al. (2017) selected 20% of the most starvation resistant flies for 80 generations; Griffin et al. (2017) selected the 10% most desiccation resistant flies for 20 generations; (Castro et al., 2018) selected the 20% mice with the longest legs for more than 20 generations; This work and previous works suggest that such strong selection likely results in a suboptimal power to identify QTNs (Kessner and Novembre, 2015). We speculate that the main motivation for choosing strong selection is the concern about an insufficient response to selection of the QTNs with weak selection. With a weak response to selection it may not be possible to distinguish the QTNs from noise generated by genetic drift. We however show that hitchhikers generated by strong selection may also lead to a substantial amount of noise, thus reducing the power to identify QTNs. We thus argue that an ideal selection regime needs to strike a balance between too strong and too weak selection. This also explains why the performance of E&R studies is very sensitive to the selection regime, where a suboptimal regime may result in a dramatically reduced power to identify the QTNs. The 90 → 20 increasing regime seems to provide a good compromise between the two opposing sources of noise, drift and hitchhiking, over a wide range of parameters.

Finally, we found that E&R studies may have a higher power to identify QTNs than GWAS. Ideally the performance of these two approaches would be compared at an identical effort, in terms of sequencing, phenotyping and time required. This is however difficult due to several reasons. In terms of sequencing an E&R study clearly requires less effort than a GWAS. Even for a powerful E&R study involving 10 replicates, solely 20 sequencing samples are necessary, whereas several hundreds (or thousands) sequencing samples are necessary for a powerful GWAS. However for GWAS with a reference panel sequencing of each strain is solely performed once and many GWAS using different traits may be performed (Schlötterer et al., 2015; Mackay et al., 2012). Thus after an initial investment, all further GWAS carried out with the reference panel will not incur additional sequencing cost. Also the phenotyping effort is difficult to compare. For example, the low-budget E&R study (*N* = 500, 5 replicates, 45 generations) had a similar performance than a GWAS with about 1500 individuals. Hence, 112, 500 (500 * 5 *45) individuals need to be phenotyped with the E&R study to achieve a similar performance than a GWAS with 1500 phenotyped individuals. In case all individuals need to be phenotyped separately GWAS thus clearly requires less phenotyping effort than an E&R study (unless GWAS is performed with a reference panel where each strain may be phenotyped multiple times for the same trait). However, with E&R studies shortcuts, that allow bulked phenotyping of populations, are frequently used. For example Turner et al. (2011) selected for large flies using a sieving apparatus.

Griffin et al. (2017) selected for desiccation resistance by exposing fly populations to dry conditions until 90% of the flies died. Hardy et al. (2017) selected for starvation resistance by depriving flies of food until 80-90% died. Other examples of bulked phenotyping, that could be used in E&R studies, are selection for flight speed using wind tunnels (Weber, 1996) and selection for pathogen resistance by breeding survivors of infections (Kraaijeveld and Godfray, 2008). In terms of our previous example, only 225 (5 * 45) bulked phenotypings of populations need to be performed for the E&R study compared to 1500 phenotypings for the GWAS. When bulked phenotyping is feasible an E&R study may thus require less phenotyping effort than a GWAS. However many traits not amenable to bulked phenotyping, like pigmentation in Drosophila, may only become accessible to E&R studies when phenotyping can be automated, e.g. with devices such as the FlySorter (Zucker and Zucker, 2017). An E&R study is usually much more time-consuming than a GWAS, as E&R typically requires multiple generations of selection. This may take several years, dependent on the used organism. An exception is a GWAS with a reference panel, where establishment of the highly inbred lines requires many generations. For example the Drosophila Genetic Reference panel (DGRP) was inbred for 20 generations (Mackay et al., 2012). This is however again a one-time investment, that is paid of by every GWAS performed with the reference panel.

For these reasons it is difficult to compare the required effort between GWAS and E&R. As a very rough guide to feasibility of an experiment, we may ask which experimental designs have been used so far. For example in Drosophila powerful E&R studies were already used: Barghi et al. (2019) used 10 replicates, *N* = 1250 and 68 generations of adaptation. To our knowledge the largest GWAS in Drosophila used several 100 individuals ((Mackay et al., 2012); not considering Pool-GWAS (Bastide et al., 2013)). Solely based on the so far used experimental designs, in Drosophila E&R may thus be a more powerful approach to identify QTNs than GWAS. On the other hand GWAS involving several thousands of individuals are regularly performed in humans (Visscher et al., 2017) where E&R is not feasible. The optimal approach for identifying QTNs will thus depend on the organism and the phenotype (e.g. if bulked phenotyping is feasible).

Our results suggest that E&R studies suffer to a lesser extent from some problems of GWAS, like the weak performance with rare alleles and alleles of small effect. With an optimal selection regime E&R studies allow to identify many rare alleles and weak effect loci. However, E&R studies suffer from own limitations. Most notably E&R studies have difficulties identifying alleles starting at a high frequency and E&R studies are highly sensitive to the selection regime. A suboptimal regime may result in a dramatically reduced power to identify QTNs, which makes E&R a more risky approach than GWAS. Moreover, with E&R studies it is not feasible to estimate the fraction of the genetic variation explained by the identified QTNs, a crucial benchmark for GWAS (Yang et al., 2011; Segura et al., 2012). Finally, due to the requirement for large populations and many generations of selection E&R will only be an option for small organism with a short generation time such as yeast, Drosophila and Caenorhabditis (Long et al., 2015). Therefore we do not view E&R as an alternative to GWAS but rather as a complementary approach with own strength and weaknesses.

## Materials and Methods

### Forward simulations

All simulations were performed with the software MimicrEE2 (Vlachos and Kofler, 2018). Briefly, MimicrEE2 allows to perform genome-wide forward simulations of evolving populations. It uses non-overlapping generations and supports simulation of temporally varying truncating selection, i.e. different numbers of individuals may be selected at each generation (Vlachos and Kofler, 2018). As not-evolved base population we obtained haplotypes that capture the pattern of natural variation of a *D. melanogaster* population from Vienna (2010) (Bastide et al., 2013). We used the recombination rate estimates for *D. melanogaster* of Comeron et al. (2012). Recombination rate estimates were obtained for 100*kb* windows from the RRC webpage (Version 2.3) (Fiston-Lavier et al., 2010). Simulations were performed for chromosomes 2L, 2R, 3L and 3R. Hence, sex chromosomes were excluded. Low recombining regions, including the entire chromosome 4, were excluded from the analysis, as these regions inflate the false positive rate (Kofler and Schlootterer, 2014). *De novo* mutations were not considered since we are mostly interested in adaptation from standing genetic variation. We simulated populations of hermaphrodites. Because males do not recombine in *D. melanogaster* we divided the recombination rate estimates by two.

If not mentioned otherwise we simulated an E&R study with a population size of *N* = 1000, 10 replicates, and 90 generations of selection. We randomly picked 100 QTNs where effect sizes were drawn from a gamma distribution with shape = 0.42. Only QTNs with allele frequencies between 5% and 95% were considered. The QTN effects were additive (no dominance or epistasis was simulated). All simulations were repeated 10 times with independent sets of randomly drawn QTNs.

### Statistical analysis

We used the CMH test (Landis et al., 1978) implemented in PoPoolation2 (Kofler et al., 2011) to identify selected loci. Previous works showed that the cmh-test has a high power to identify selected loci in E&R studies (Vlachos and Kofler, 2018; Kofler and Schlötterer, 2014). The CMH test is based on a meta-analysis of a 2*2\**k* contingency table. This contingency table contains for each replicate (k) the counts of the major and the minor allele (2), for the base and the evolved population (2). The null hypothesis is the absence of differentiation between base and evolved populations. Additionally, we evaluated the performance of two time-series bases approaches: CLEAR and an adaptation of the CMH test to E&R studies (Iranmehr et al., 2017; Spitzer et al., 2019). To obtain time-series data we performed simulations with the default parameters, requesting an output each 10^*th*^ generations (10 time points in total). We provided the harmonic mean of the population size (i.e. the number of selected individuals) as estimate of *Ne* required by the adapted CMH test. As CLEAR is very slow, we solely analyzed the data for a single chromosome arm (2L). Finally, we used the programming language R (R Core Team, 2014) and the library ROCR to generate ROC curves and to compute the area under the ROC curve (AUC) (Sing et al., 2005).

### Maximizing the phenotypic response

Truncating selection may either be performed to identify the QTNs or to maximize the phenotypic response. To compare the performance of selection regimes that are best suited for these two tasks we simulated truncating selection for 90 generations. The selection regime with the highest power to identify QTNs was identified as describe above (E&R study with *N* = 1000, 10 replicates; 10 independent sets of QTNs). To identify the regime which maximizes the phenotypic value we computed the response to selection (*R*: phenotypic difference between evolved and base population) for all increasing regimes using steps of 10% selected individuals (E&R study with *N* = 1000, 1 replicate; 100 independent sets of QTNs). Finally we picked the regime with the largest average response.

### GWAS

All GWAS were performed with the software SNPtest (Marchini et al., 2007). We used an additive model and raw phenotypic data for a quantitative trait (parameters: -frequentist 1 -method expected - use_raw_phenotypes). All simulations were performed with 10 independent sets of SNPs.

## Acknowledgements

We thank Christian Schlötterer, Nick Barton and Ilse Höllinger for comments, and Rupert Mazzucco for help with the computer cluster. The results have been partly obtained using the Vienna Scientific Cluster (VSC). We thank all members of the Institute of Population Genetics for feedback and support This work was supported by an Austrian Science Foundation (FWF) grant (P29016-B25) to RK.

## References

J. G. Baldwin-Brown, A. D. Long, and K. R. Thornton. The Power to Detect Quantitative Trait Loci Using Resequenced, Experimentally Evolved Populations of Diploid, Sexual Organisms. Molecular biology and evolution, 31:1040–55, 2014.

N. Barghi, V. Nolte, T. Taus, A. M. Jakšić, R. Tobler, F. Mallard, M. Dolezal, C. Schlötterer, R. Kofler, and K. A. Otte. Genetic redundancy fuels polygenic adaptation in *Drosophila*. PLOS Biology, 17(2):e3000128, 2019.

N. H. Barton, A. M. Etheridge, and A. Véber. The infinitesimal model: Definition, derivation, and implications. Theoretical Population Biology, 118:50–73, 2017.

H. Bastide, A. Betancourt, V. Nolte, R. Tobler, P. Stöbe, A. Futschik, and C. Schlötterer. A Genome-Wide, Fine-Scale Map of Natural Pigmentation Variation in *Drosophila melanogaster*. PLoS Genetics, 9: e1003534, 2013.

J. Castro, M. N. Yancoskie, M. Marchini, S. Belohlavy, W. H. Beluch, R. Naumann, I. Skuplik, J. Cobb, H. Nick, C. Rolian, and Y. F. Chan. An integrative genomic analysis of the Longshanks selection experiment for longer limbs in mice. bioRxiv, 2018.

J. Comeron, R. Ratnappan, and S. Bailin. The Many Landscapes of Recombination in *Drosophila melanogaster*. PLoS Genetics, 8(10):e1002905, 2012.

E. L. Dittmar, C. G. Oakley, J. K. Conner, B. A. Gould, and D. W. Schemske. Factors influencing the effect size distribution of adaptive substitutions. Proceedings of the Royal Society B: Biological Sciences, 283: 20153065, 2016.

J. W. Dudley and R. J. Lambert. 100 Generations of Selection for Oil and Protein in Corn, volume 24. 2010.

M. El-Soda, M. Malosetti, B. J. Zwaan, M. Koornneef, and M. G. Aarts. Genotype - environment interaction QTL mapping in plants: Lessons from *Arabidopsis*. Trends in Plant Science, 19(6):390–398, 2014.

D. Falconer. Early Selection Experiments. Annual Review of Genetics, 26(1):1–14, 1992.

D. Falconer and T. Mackay. Introduction to quantitative genetics. Harlow: Pearson, Prentice Hall., 1960.

R. Fisher. The genetical theory of natural selection. Oxford Univ. Press, Oxford, 1930.

A.-S. Fiston-Lavier, N. D. Singh, M. Lipatov, and D. A. Petrov. *Drosophila melanogaster* recombination rate calculator. Gene, 463(1-2):18–20, 2010.

T. Garland and M. R. Rose. Experimental evolution: concepts, methods, and applications of selection experiments. University of California Press Berkeley, CA, 2009.

G. Gibson. Rare and common variants: twenty arguments. Nature reviews Genetics, 13(2):135–45, 2011.

G. Gibson. Population genetics and GWAS: A primer. PLOS Biology, 16:e2005485, 2018.

J. L. Graves, K. L. Hertweck, M. A. Phillips, M. V. Han, L. G. Cabral, T. T. Barter, L. F. Greer, M. K. Burke, L. D. Mueller, M. R. Rose, and N. Singh. Genomics of parallel experimental evolution in *Drosophila*. Molecular Biology and Evolution, 34(4):831–842, 2017.

P. C. Griffin, S. B. Hangartner, A. Fournier-Level, and A. A. Hoffmann. Genomic Trajectories to Desiccation Resistance: Convergence and Divergence Among Replicate Selected *Drosophila* Lines. Genetics, 205(2): 871–890, 2017.

C. Hardy, M. Burke, L. Everett, M. Han, K. Lantz, and A. Gibbs. Genome-Wide Analysis of Starvation-Selected *Drosophila melanogaster*—A Genetic Model of Obesity. Molecular Biology and Evolution, 35: 50–65, 2017.

T. Hastie, R. Tibshirani, and J. Friedman. The Elements of Statistical Learning: Data Mining, Inference, and Prediction. Springer series in statistics, Springer-Verlag New York, 2009.

B. Hayes and M. E. Goddard. The distribution of the effects of genes affecting quantitative traits in livestock. Genetics, selection, evolution, 33(3):209–229, 2001.

A. Iranmehr, A. Akbari, C. Schlötterer, and V. Bafna. CLEAR: Composition of likelihoods for evolve and resequence experiments. Genetics, 206(2):1011–1023, 2017.

P. D. Keightley and G. Bulfield. Detection of quantitative trait loci from frequency changes of marker alleles under selection. Genetical Research, 62:195, 1993.

J. K. Kelly and K. A. Hughes. Pervasive Linked Selection and Intermediate-Frequency Alleles Are Implicated in an Evolve-and-Resequencing Experiment of *Drosophila simulans*. Genetics, page genetics.301824.2018, 2018.

D. Kessner and J. Novembre. Power analysis of artificial selection experiments using efficient whole genome simulation of quantitative traits. Genetics, 199(4):991–1005, 2015.

R. Kofler and C. Schlötterer. A Guide for the Design of Evolve and Resequencing Studies. Molecular biology and evolution, 31(2):474–483, 2014.

R. Kofler, R. V. Pandey, and C. Schlötterer. PoPoolation2: identifying differentiation between populations using sequencing of pooled DNA samples (Pool-Seq). Bioinformatics, 27:3435–3436, 2011.

A. Korte and A. Farlow. The advantages and limitations of trait analysis with GWAS: a review. Plant methods, 9(1):29, 2013.

K. Kosheleva and M. M. Desai. Recombination alters the dynamics of adaptation on standing variation in laboratory yeast populations. Molecular Biology and Evolution, 35:180–201, 2017.

A. Kraaijeveld and H. Godfray. Selection for resistance to a fungal pathogen in *Drosophila melanogaster*. Heredity, 100(4):400, 2008.

E. S. Lander and N. J. Schork. Genetic dissection of complex traits. Science, 265(5181):2037–48, 1994.

J. R. Landis, E. R. Heyman, and G. G. Koch. Average partial association in three-way contingency tables: a review and discussion of alternative tests. International Statistical Review/Revue Internationale de Statistique, 46(3):237–254, 1978.

A. Long, G. Liti, A. Luptak, and O. Tenaillon. Elucidating the molecular architecture of adaptation via evolve and resequence experiments. Nature Reviews Genetics, 16(10):567–582, 2015.

J. B. Losos, S. J. Arnold, G. Bejerano, E. D. Brodie III, D. Hibbett, H. E. Hoekstra, D. P. Mindell, A. Monteiro, C. Moritz, H. A. Orr, D. A. Petrov, S. S. Renner, R. E. Ricklefs, P. S. Soltis, and T. L. Turner. Evolutionary biology for the 21st century. PLoS Biology, 11(1):e1001466, 2013.

T. F. Mackay. The genetic architecture of quantitative traits. Annual review of genetics, 35:303–39, 2001.

T. F. C. Mackay, E. A. Stone, and J. F. Ayroles. The genetics of quantitative traits: challenges and prospects. Nature reviews. Genetics, 10(8):565–77, 2009.

T. F. C. Mackay, S. Richards, E. A. Stone, A. Barbadilla, J. F. Ayroles, D. Zhu, S. Casillas, Y. Han, M. M. Magwire, J. M. Cridland, M. F. Richardson, R. R. H. Anholt, M. Barrón, C. Bess, K. P. Blankenburg, M. A. Carbone, D. Castellano, L. Chaboub, L. Duncan, Z. Harris, M. Javaid, J. C. Jayaseelan, S. N. Jhangiani, K. W. Jordan, F. Lara, F. Lawrence, S. L. Lee, P. Librado, R. S. Linheiro, R. F. Lyman, A. J. Mackey, M. Munidasa, D. M. Muzny, L. Nazareth, I. Newsham, L. Perales, L.-L. Pu, C. Qu, M. Ràmia, J. G. Reid, S. M. Rollmann, J. Rozas, N. Saada, L. Turlapati, K. C. Worley, Y.-Q. Wu, A. Yamamoto, Y. Zhu, C. M. Bergman, K. R. Thornton, D. Mittelman, and R. A. Gibbs. The *Drosophila melanogaster* Genetic Reference Panel. Nature, 482(7384):173–8, 2012.

J. Marchini, L. Cardon, M. Phillips, and P. Donnelly. The effects of human population structure on large genetic association studies. Nature Genetics, 36:512–517, 2004.

J. Marchini, S. Howie, B.and Myers, G. McVean, and P. Donnelly. A new multipoint method for genome-wide association studies by imputation of genotypes. Nature Genetics, 39:906–913, 2007.

N. E. Martins, V. G. Faria, V. Nolte, C. Schlötterer, L. Teixeira, E. Sucena, and S. Magalhães. Host adaptation to viruses relies on few genes with different cross-resistance properties. Proceedings of the National Academy of Sciences, 111(43):15597–15597, 2014.

T. H. E. Meuwissen, B. J. Hayes, and M. E. Goddard. Prediction of Total Genetic Value Using Genome-Wide Dense Marker Maps. Genetics, 157(4):1819–1829, 2001.

P. Orozco-Terwengel, M. Kapun, V. Nolte, R. Kofler, T. Flatt, and C. Schlötterer. Adaptation of *Drosophila* to a novel laboratory environment reveals temporally heterogeneous trajectories of selected alleles. Molecular Ecology, 21(20):4931–4941, 2012.

M. A. Phillips, G. A. Rutledge, J. N. Kezos, Z. S. Greenspan, S. Matty, H. Arain, L. D. Mueller, A. Talbott, M. R. Rose, and P. Shahrestani. Effects of evolutionary history on genome wide and phenotypic convergence in *Drosophila* populations. BMC Genomics, 19(1):1–17, 2018.

R Core Team. R: A Language and Environment for Statistical Computing. R Foundation for Statistical Computing, Vienna, Austria, 2014. URL http://www.R-project.org/.

S. Ripke, B. M. Neale, A. Corvin, J. T. Walters, K. H. Farh, P. A. Holmans, P. Lee, B. Bulik-Sullivan, D. A. Collier, H. Huang, and et al. Biological insights from 108 schizophrenia-associated genetic loci. Nature, 511(7510):421–427, 2014.

A. Robertson. A Theory of Limits in Artificial Selection. Genetics, 144(95):234–249, 1960.

A. Robertson. Some optimum problems in individual selection. Theoretical Population Biology, 1(1):120–127, 1970a.

A. Robertson. A theory of limits in artificial selection with many linked loci. In Mathematical topics in population genetics, pages 246–288. Springer, 1970b.

M. Rockman. THE QTN PROGRAM AND THE ALLELES THAT MATTER FOR EVOLUTION: ALL THAT’S GOLD DOES NOT GLITTER. Evolution, 66(1):1–17, 2011.

C. Schlötterer, R. Tobler, R. Kofler, and V. Nolte. Sequencing pools of individuals-mining genome-wide polymorphism data without big funding. Nature Reviews Genetics, 15(11):749–763, 2014.

C. Schlötterer, R. Kofler, E. Versace, R. Tobler, and S. Franssen. Combining experimental evolution with next-generation sequencing: a powerful tool to study adaptation from standing genetic variation. Heredity, 114(5):431–440, 2015.

V. Segura, B. J. Vilhjálmsson, A. Platt, A. Korte, U. Seren, Q. Long, and M. Nordborg. An efficient multi-locus mixed-model approach for genome-wide association studies in structured populations. Nature Genetics, 44(7):825–830, 2012.

T. Sing, O. Sander, N. Beerenwinkel, and T. Lengauer. ROCR: visualizing classifier performance in R. Bioinformatics, 21:3940–3941, 2005.

K. Spitzer, M. Pelizzola, and A. Futschik. Modifying the Chi-square and the CMH test for population genetic inference: adapting to over-dispersion. 2019. URL http://arxiv.org/abs/1902.08127.

J. Stapley, J. Reger, P. G. D. Feulner, C. Smadja, J. Galindo, R. Ekblom, C. Bennison, A. D. Ball, A. P. Beckerman, and J. Slate. Adaptation genomics: the next generation. Trends in Ecology & Evolution, 25 (12):705–712, 2010.

H. Teotonio, S. Carvalho, D. Manoel, M. Roque, and I. M. Chelo. Evolution of Outcrossing in Experimental Populations of *Caenorhabditis elegans*. PLoS ONE, 7:e35811, 2012.

R. Tobler, S. U. Franssen, R. Kofler, P. Orozco-Terwengel, V. Nolte, J. Hermisson, and C. Schlötterer. Massive habitat-specific genomic response in *D. melanogaster* populations during experimental evolution in hot and cold environments. Molecular Biology and Evolution, 31(2):364–375, 2014.

H. Topa, Á. Jónás, R. Kofler, C. Kosiol, and A. Honkela. Gaussian process test for high-throughput sequencing time series: Application to experimental evolution. Bioinformatics, 31(11):1762–1770, 2015.

T. Turner and P. Miller. Investigating Natural Variation in *Drosophila* Courtship Song by the Evolve and Resequence Approach. Genetics, 191:633–642, 2012. doi: {10.1534/genetics.112.139337}.

T. Turner, A. D., S. Andrew, T. Fields, W. R. Rice, and A. M. Tarone. Population-Based Resequencing of Experimentally Evolved Populations Reveals the Genetic Basis of Body Size Variation in *Drosophila melanogaster*. PLoS Genetics, 7(3):e1001336, 2011. doi: {10.1371/journal.pgen.1001336}.

P. M. Visscher, W. G. Hill, and N. R. Wray. Heritability in the genomics era — concepts and misconceptions. Nature Reviews Genetics, 9(4):255–266, 2008.

P. M. Visscher, M. A. Brown, M. I. McCarthy, and J. Yang. Five years of GWAS discovery. American journal of human genetics, 90(1):7–24, 2012.

P. M. Visscher, N. R. Wray, Q. Zhang, P. Sklar, M. I. McCarthy, M. A. Brown, and J. Yang. 10 Years of GWAS Discovery: Biology, Function, and Translation. American Journal of Human Genetics, 101(1): 5–22, 2017.

C. Vlachos and R. Kofler. MimicrEE2: Genome-wide forward simulations of Evolve and Resequencing studies. PLoS Computational Biology, 14:e1006413, 2018.

A. M. Wannier, Timothy M. and Kunjapur, D. P. Rice, M. J. McDonald, M. M. Desai, and G. M. Church. Adaptive evolution of genomically recoded *Escherichia coli*. Proceedings of the National Academy of Sciences, 115:3090–3095, 2018.

K. E. Weber. Large genetic change at small fitness cost in large populations of *Drosophila melanogaster* selected for wind tunnel flight: Rethinking fitness surfaces. Genetics, 144(1):205–213, 1996.

J. Yang, S. H. Lee, M. E. Goddard, and P. M. Visscher. GCTA: A tool for genome-wide complex trait analysis. American Journal of Human Genetics, 88(1):76–82, 2011.

D. S. Zucker and M. A. Zucker. Insect singulating device, 2017. US Patent App. 14/851,610.

