## supplement for "Optimizing the power to identify the genetic basis of complex traits with Evolve and Resequence studies"

March 20, 2019

#### 1 Supplementary figures

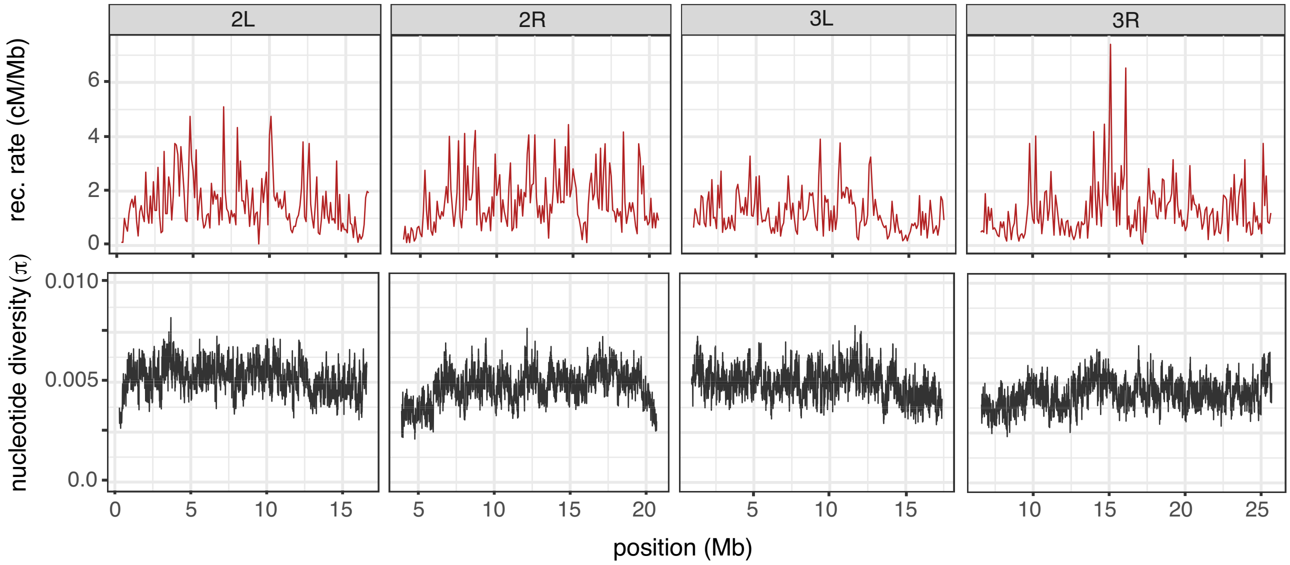


Figure 1: The genomic landscape used in the simulations. The recombination rate is from Comeron et al. (2012) and the base population captures the nucleotide diversity of a natural *D. melanogaster* population from Vienna (Bastide et al., 2013). The window size size is 100kb.

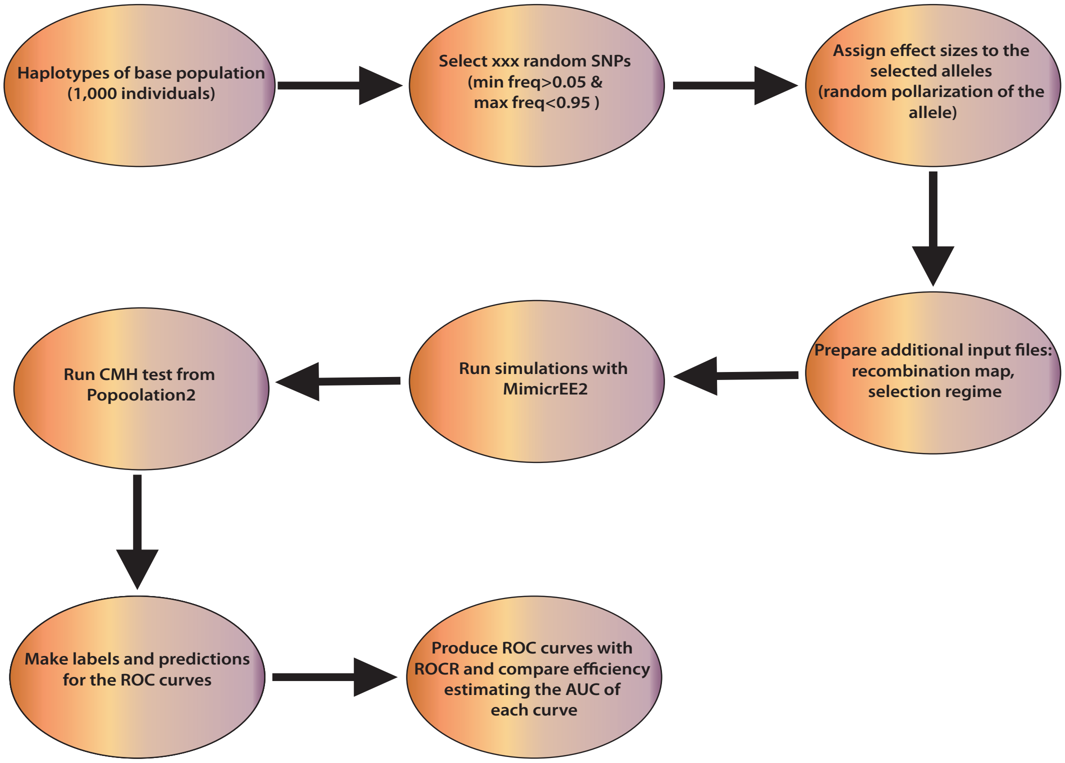


Figure 2: Overview of our simulation pipeline. The best selection regime was identified based on the largest Area Under the Curve (AUC) of each Receiver Operating Characteristic (ROC) plot.

#### 1.1 Allele frequency changes in truncation selection simulations

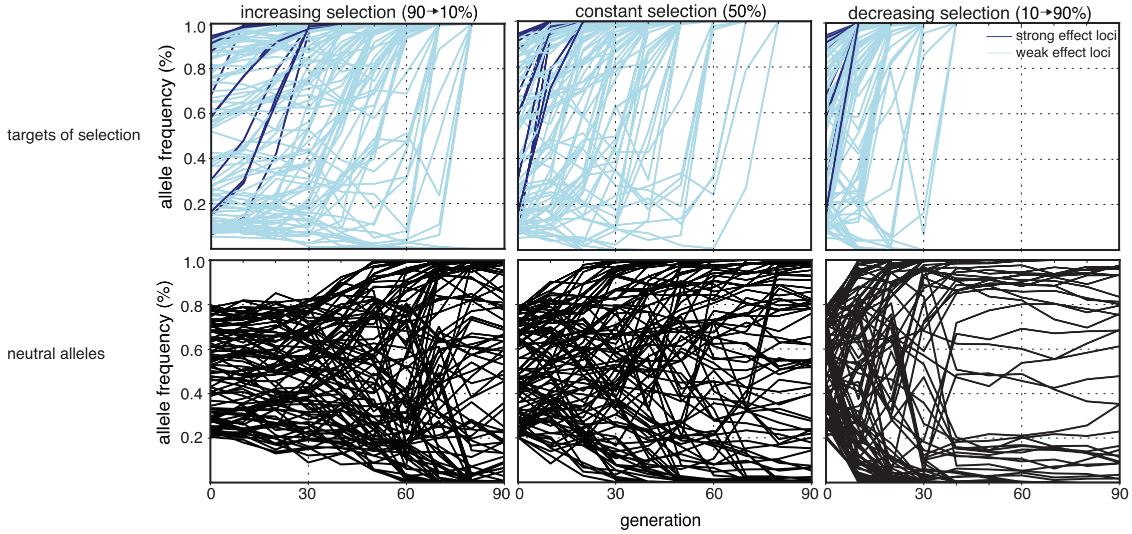


Figure 3: Trajectories of selected (blue) and not-selected alleles (black) for three different selection regimes (top). Strong effect loci (effect size >1) are shown in dark blue whereas weak effect loci (effect size $\leq 1$) are shown in light blue. We also show the trajectories of 100 randomly picked neutral alleles that have starting frequency between 0.2 and 0.8. Data are shown for one replicate. Note that fixation of strong effect loci is delayed in the increasing regime (left) compared to the constant regime (middle) and the decreasing regime (right).

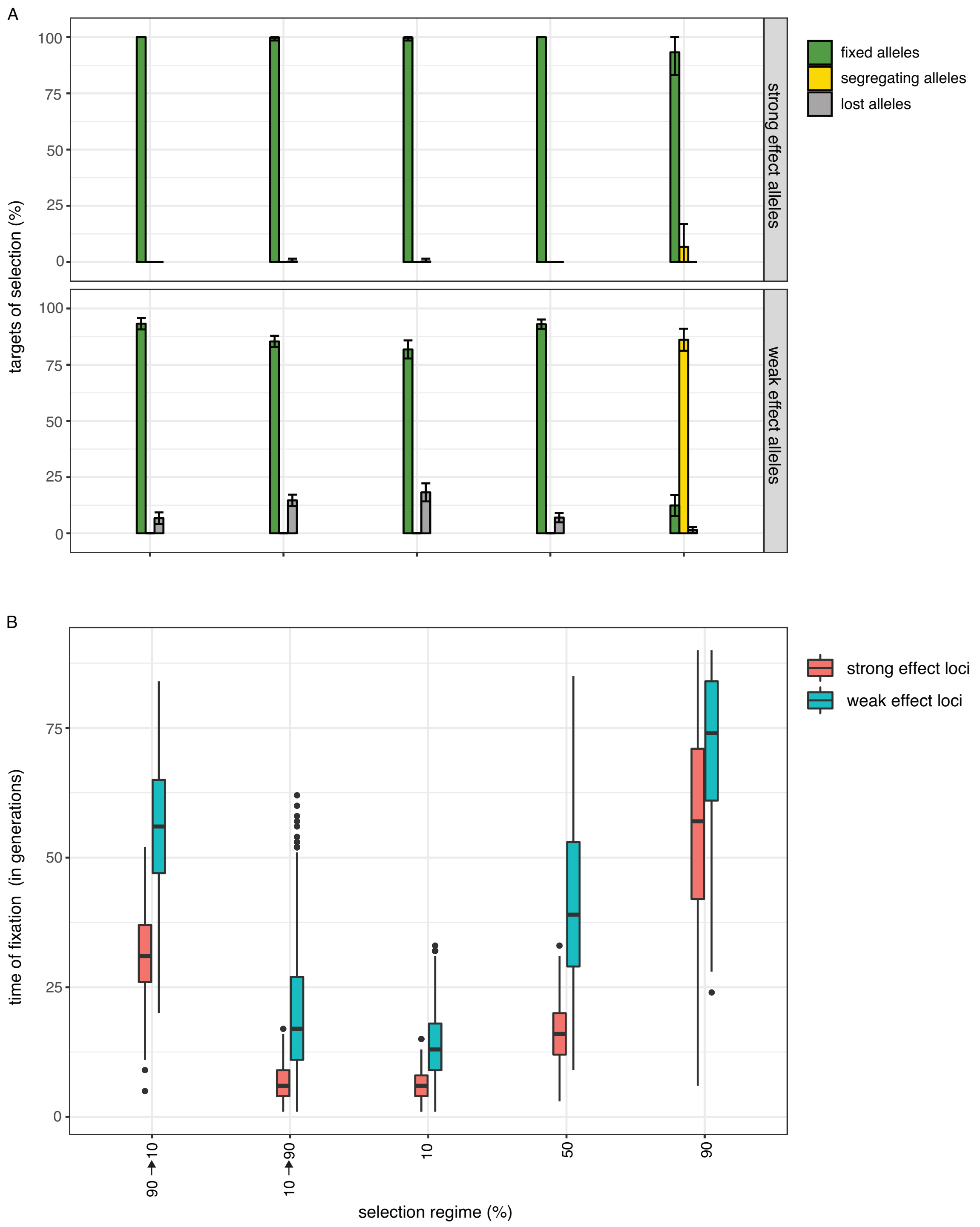


Figure 4: Fate of selected loci for 5 different selection regimes. The influence of an increasing (90 → 10%), a decreasing (10 → 90%) and three constant regimes (10%, 50%, 90%) was evaluated. A) Fraction of fixed (green), lost (grey) and segregating loci (yellow) after 90 generations of selection. Results are shown for strong and weak effect loci. B) Time to fixation for strong and weak effect loci. Note that a linearly increasing regime delays fixation of selected loci compared to the constant regime with 50% selected individuals (Wilcoxon rank sum test; $p < 0.05$).

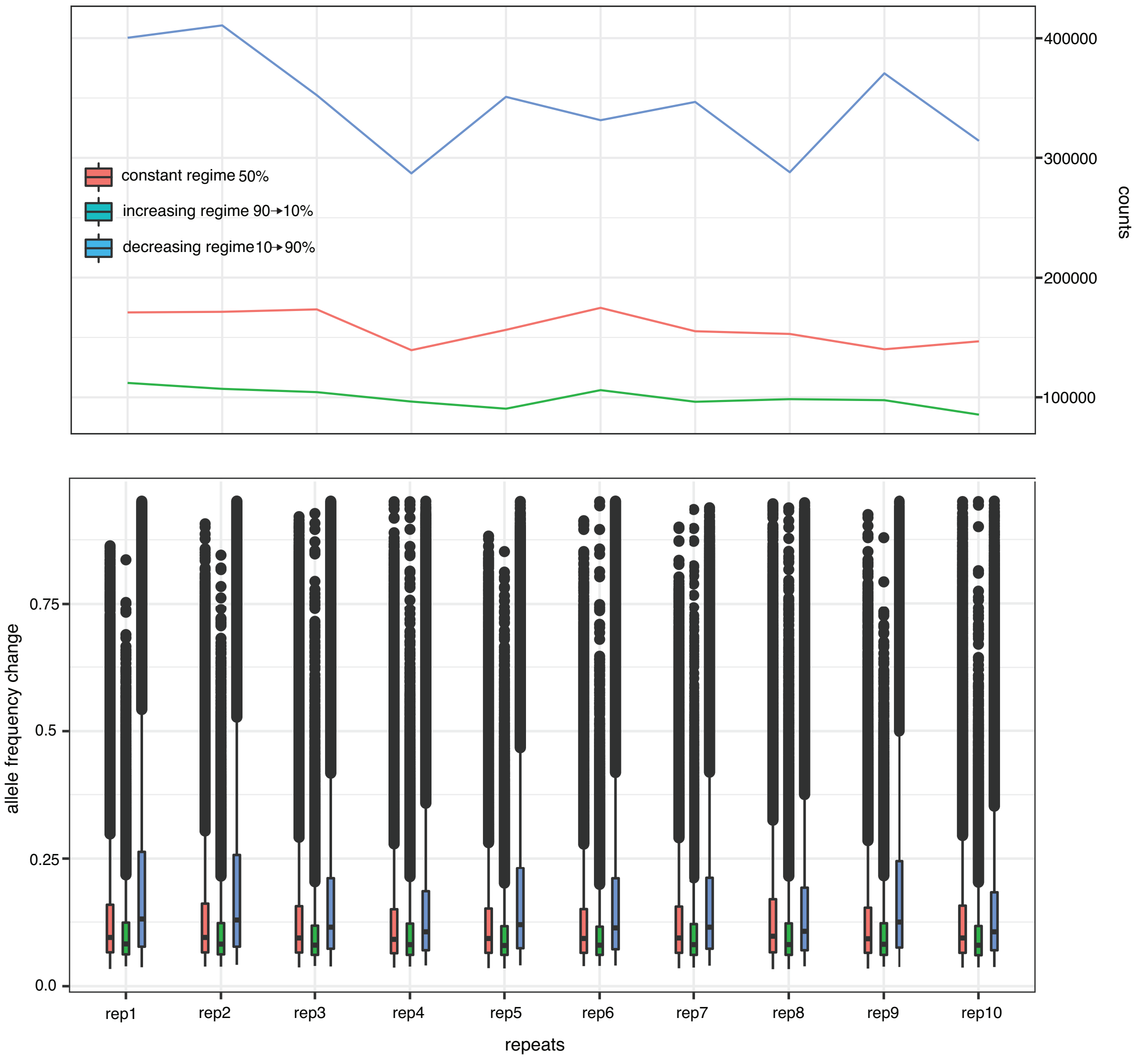


Figure 5: Overview of hitchhiking alleles for an E&R study ($N = 1000$, 10 replicates, 90 generations) using a linearly increasing, a constant and a linearly decreasing selection regime. Results are shown for 10 different sets of selected loci ($rep1..rep10$). We defined hitchhikers as loci where allele frequencies change in the same direction in all 10 replicates (excluding the selected loci). The number of identified hitchhikers is shown as line plots (top panel) and the allele frequency change of the hitchhikers as boxplots (bottom panel). Note that the linearly increasing regime consistently generates fewer hitchhikers which also show a weaker response to selection than hitchhikers from the constant and decreasing regimes (Wilcoxon rank sum test; $p < 0.05$ in each repeat).

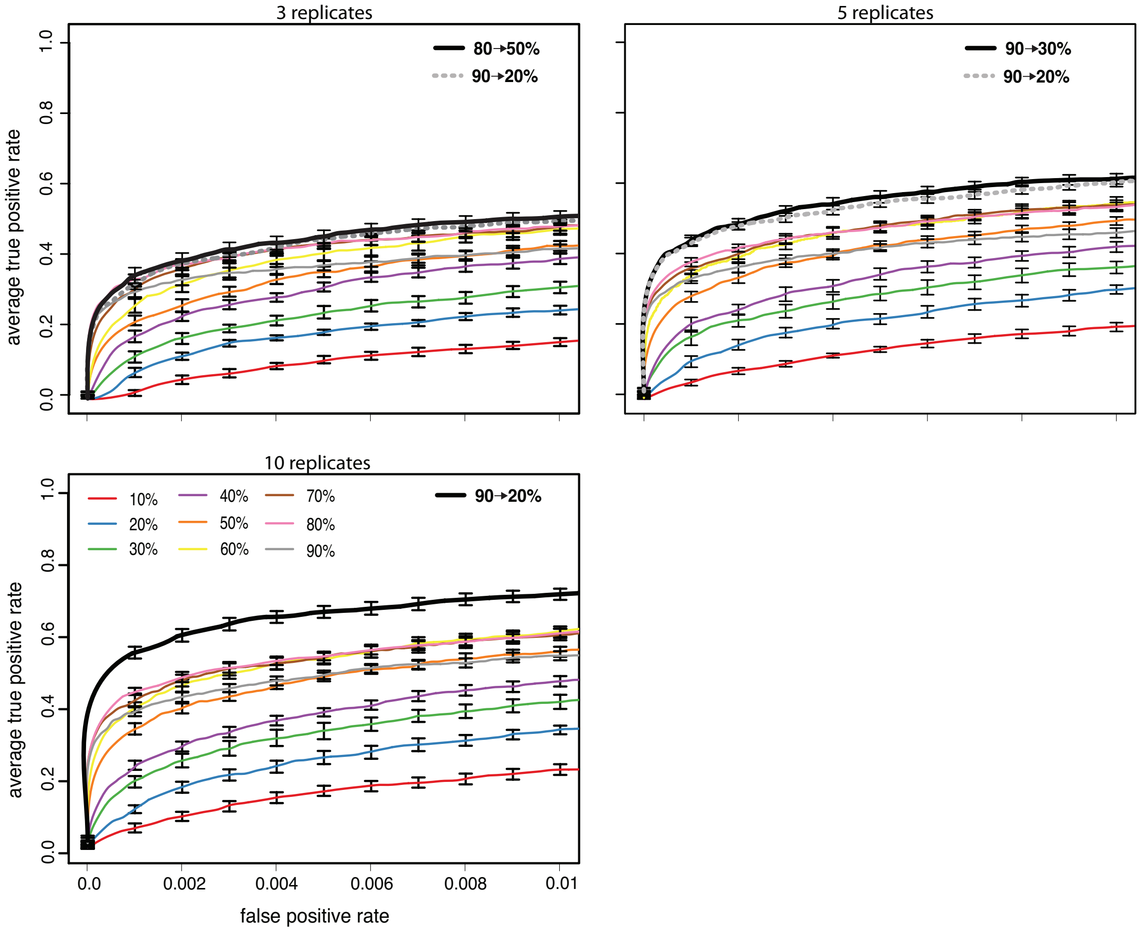


Figure 6: Influence of the number of replicates ($default = 10$). Although linearly increasing regimes consistently perform best or equally well than the constant regimes, the advantage of linearly increasing regimes is most pronounced for 5 or more replicates. (Wilcoxon rank sum test with pAUC; $p_3 = 0.5$, $p_5 = 0.0004$, $p_{10} = 0.0002$).

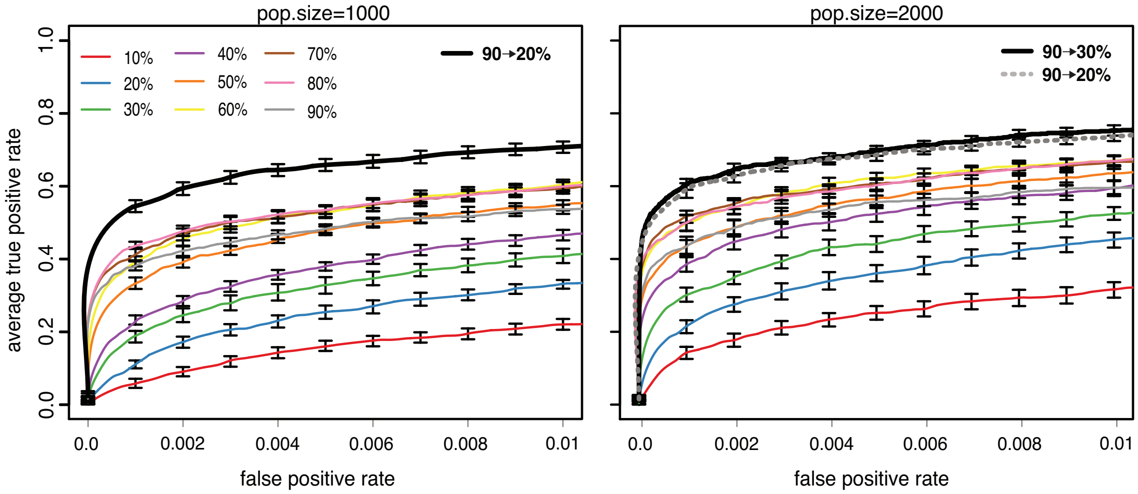


Figure 7: Influence of the population size ($default = 1000$). Linearly increasing selection regimes consistently perform better than the constant regimes (Wilcoxon rank sum test with pAUC; $p_{1000} = 0.0002$, $p_{2000} = 1e-05$).

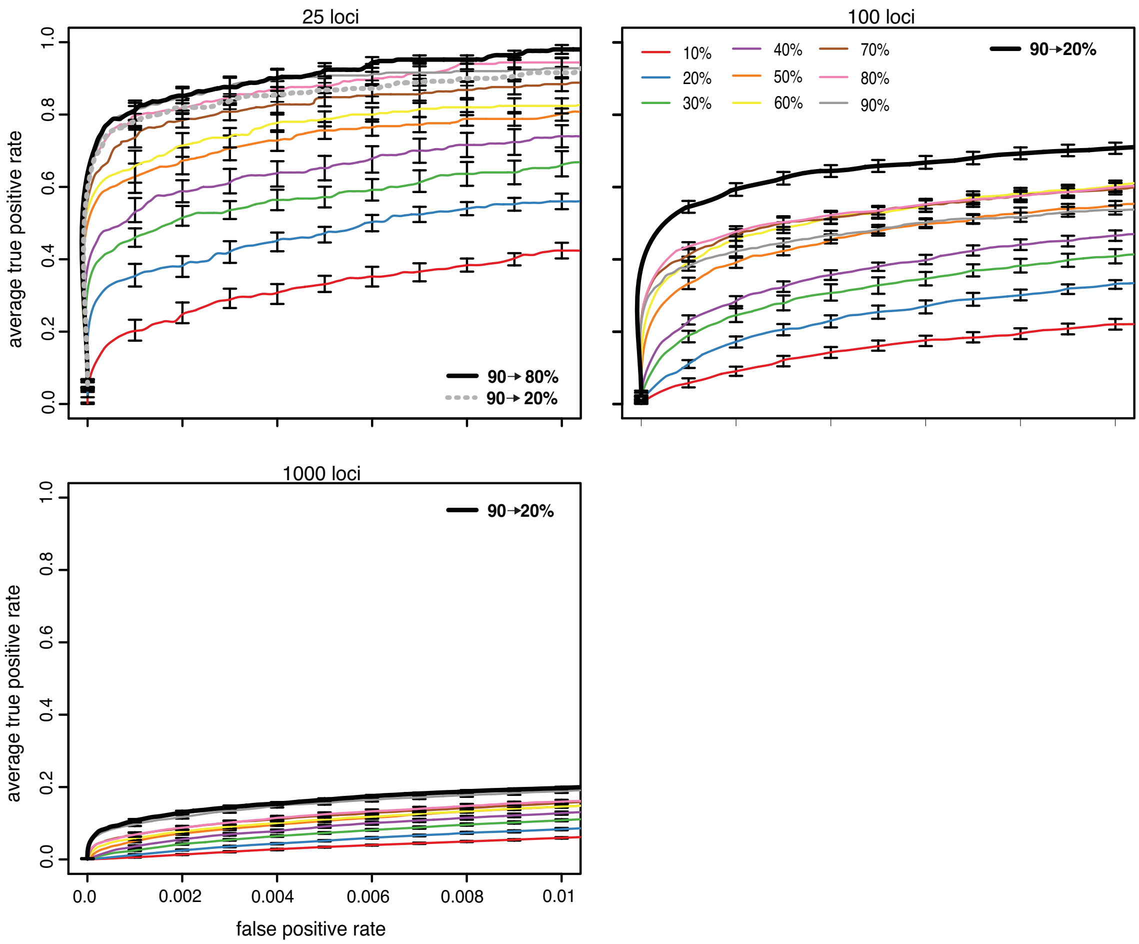


Figure 8: Influence of the number of QTNs ($default = 100$). Although linearly increasing selection regimes consistently perform best or equally well than the constant regimes, the advantage of linearly increasing regimes is most pronounced for intermediate numbers of QTNs (Wilcoxon rank sum test with pAUC; $p_{25} = 0.73$, $p_{100} = 0.0002$, $p_{1000} = 0.14$).

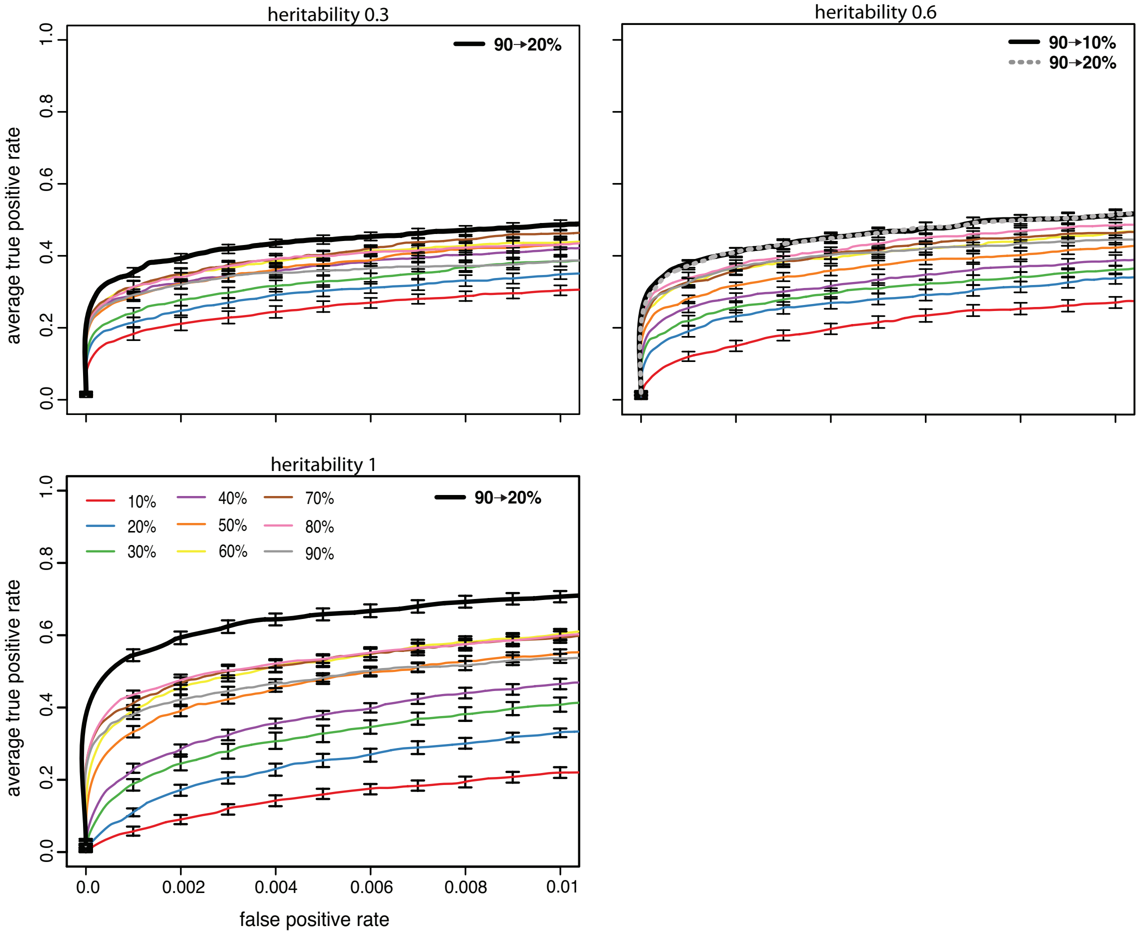


Figure 9: Influence of the heritability. The increasing selection regime consistently outperforms the constant selection regime, where the advangtage is most pronounced for high heritabilities (Wilcoxon rank sum test with pAUC; $p_{0.3} = 0.035$, $p_{0.6} = 0.035$, $p_1 = 0.0002$). Note that the influence of the selection regime diminishes with decreasing heritability (the ROC curves become more similar).

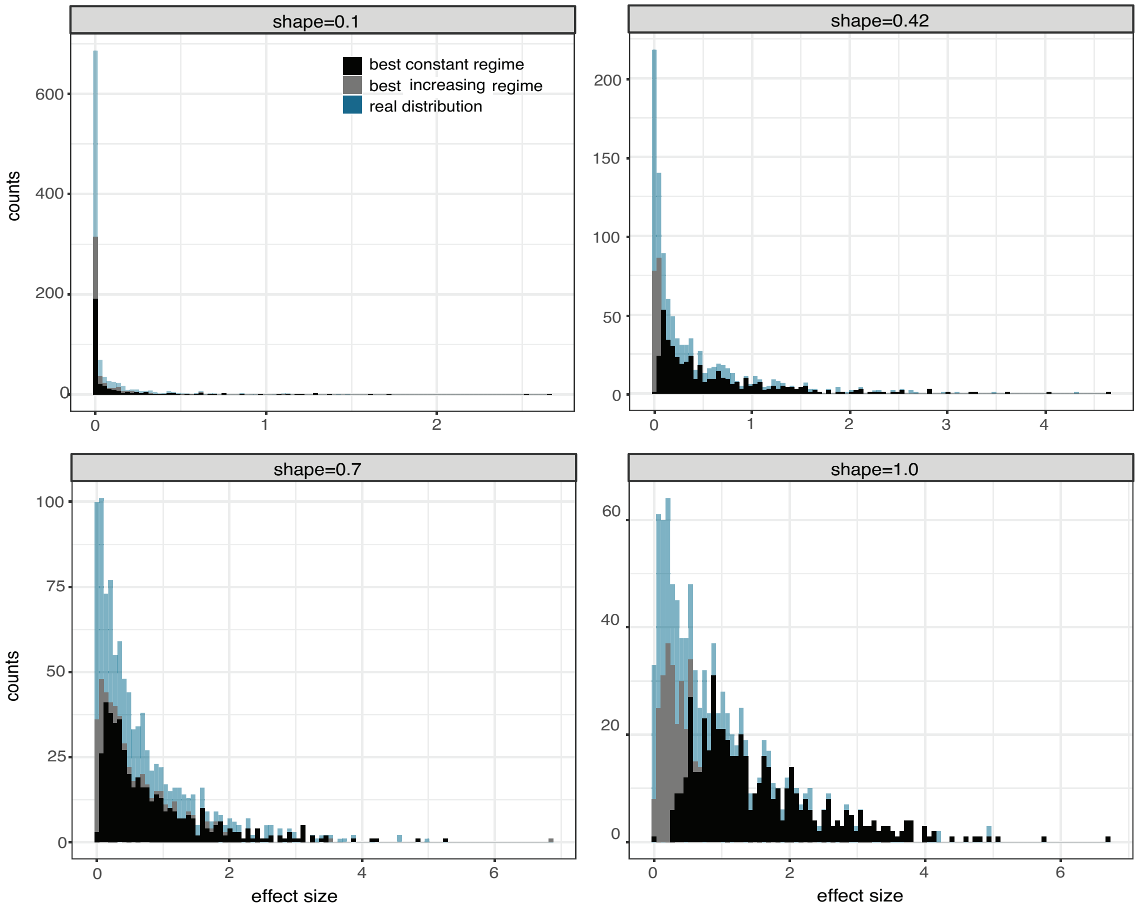


Figure 10: Histogram of the effect sizes of QTNs recovered with the best increasing (grey) and the best constant (black) regime compared to all simulated QTNs (blue). Results are shown for different gamma distributed effect sizes (top panel). Only QTNs among the 2,000 most significant SNPs were considered for this analysis. Results show the sum over 10 experiments with different random sets of QTNs. Note that the best increasing regime enables us to recover the effect size distribution more accurately than the best constant regime.

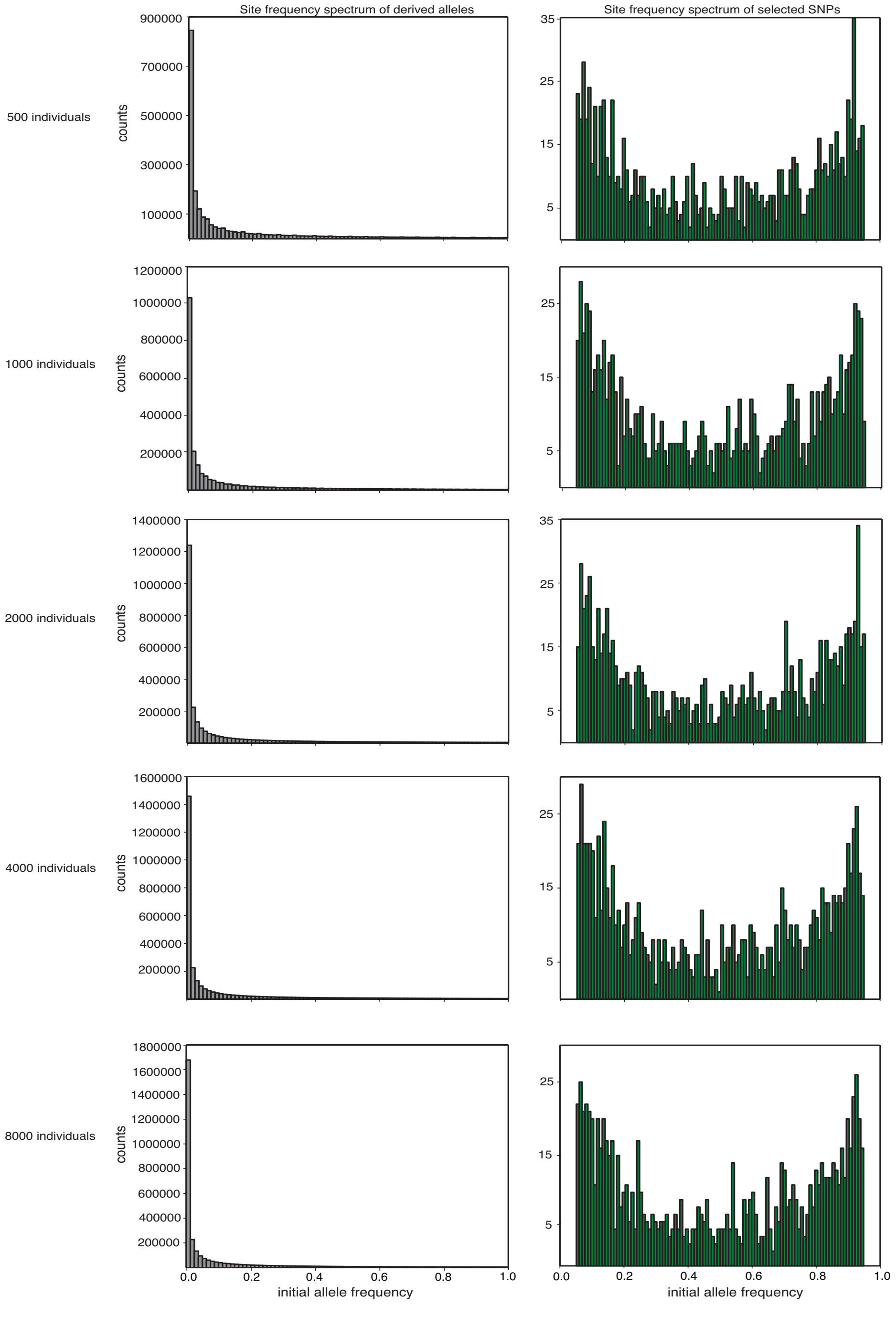


Figure 11: Site frequency spectrum of segregating SNPs (left column) and of the selected QTNs (right column). Results are shown for the five different populations sizes used in the simulated GWAS. Note that segregating SNPs were polarized by the derived allele whereas QTNs were polarized by the sign of the effect size ($+a$).

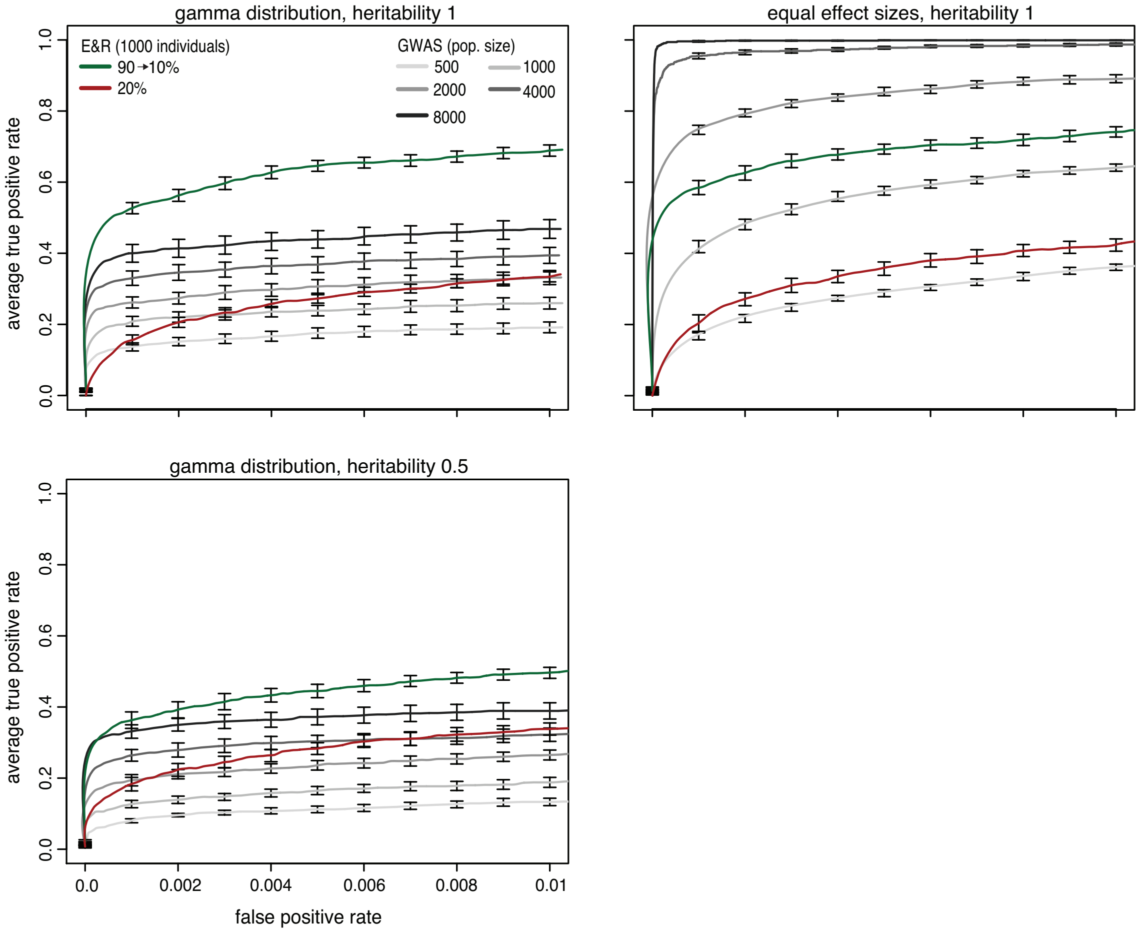


Figure 12: Influence of the heritability and effect size distribution on the performance of E&R and GWA studies. An E&R study with an increasing regime outperforms GWAS when the trait has a low heritability but not when effect sizes of QTNs are identical (Wilcoxon rank sum test with pAUC; $E\&R_{1000}$ vs $GWAS_{8000}$; with heritability 0.5: $p = 0.023$, with equal effect size: $p = 1.08e - 05$).

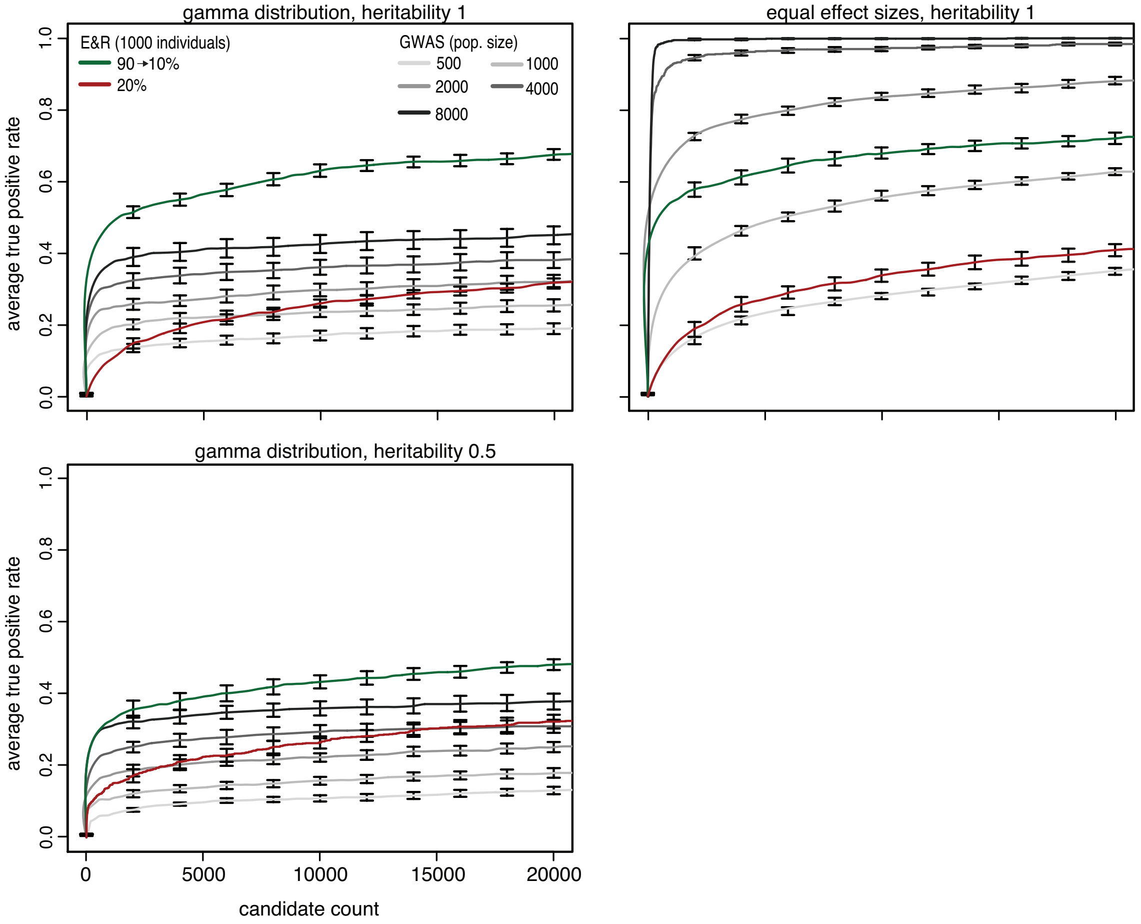


Figure 13: Influence of the heritability and effect size distribution on the performance of E&R and GWAS. The top 20,000 candidate loci are shown on the abscissa.

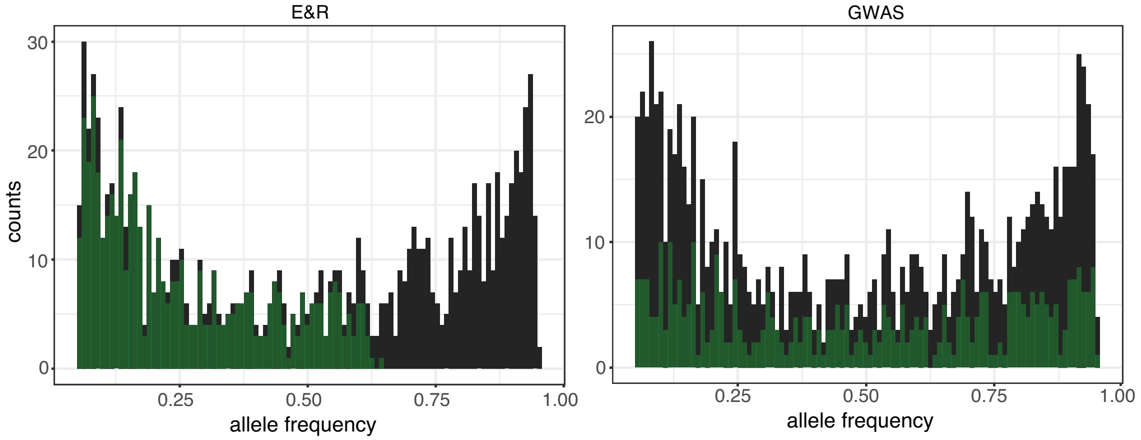


Figure 14: Starting allele frequency of QTNs identified with GWAS and E&R studies (green) compared to expectations (black). Only QTNs among the 2,000 most significant SNPs were considered for this analysis.

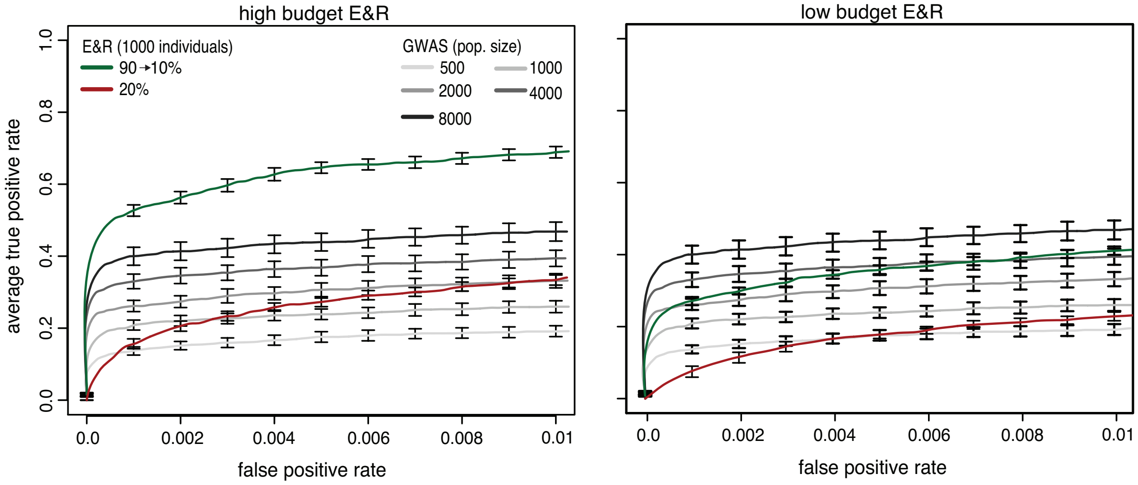


Figure 15: Performance of a low-budget E&R study ($N = 500$, 5 replicates, 45 generations), a high-budget E&R study ($N = 1000$, 10 replicates, 90 generations) and multiple GWAS with different populations sizes. A low-budget E&R study, with an improved selection regime has a similar performance than a GWAS with $1500 - 2000$ individuals.

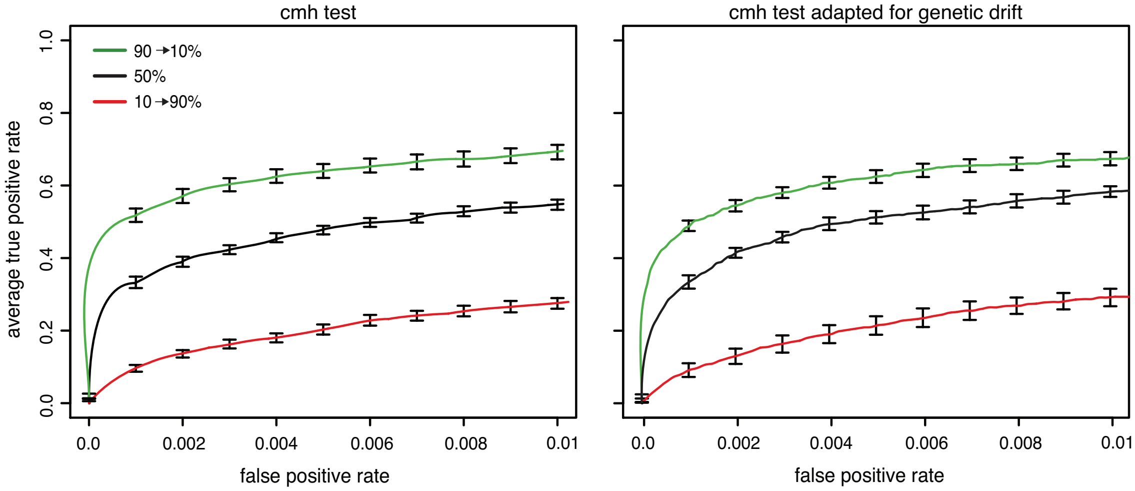


Figure 16: Performance of the classic cmh-test and a time-series based cmh-test adapted to E&R studies (Spitzer et al., 2019). For the adapted cmh-test we used allele frequency estimates at each $10^t h$ generation (10 time points in total with 90 generations of selection) and estimates of $N_e$ based on the harmonic mean of the number of selected individuals (constant regime $N_e$=500; increasing/decreasing regime $N_e$=360). With the adapted cmh-test the increasing regime performed better than the constant regime (Wilcoxon rank-sum test with pAUC; $90 \rightarrow 10\%$ vs 50%; $p = 0.0002$) and the influence of the selection regime on the performance was significant (Kruskal–Wallis rank-sum test with pAUC; $p = 4.59e - 06$).

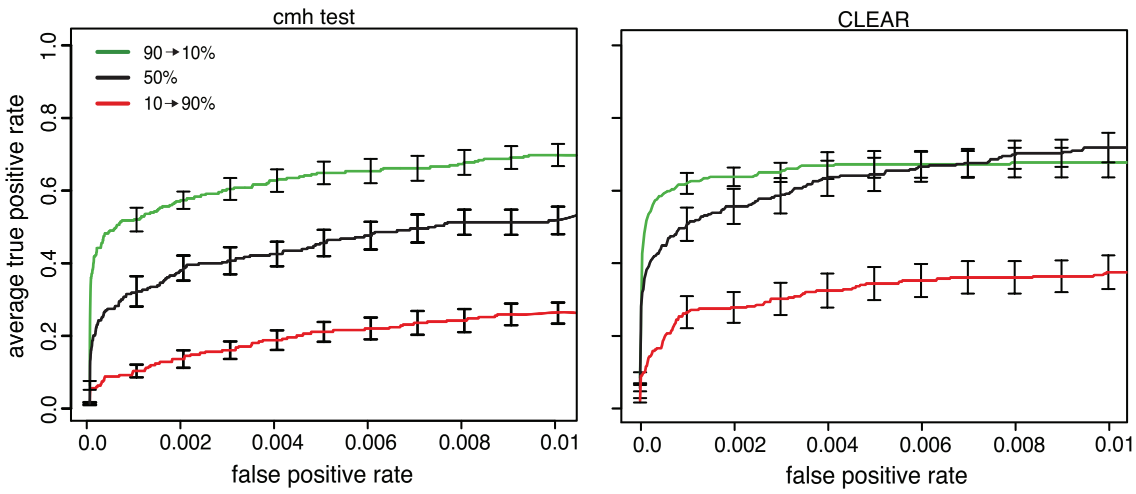


Figure 17: Performance of the cmh-test and of CLEAR, a time-series based approach for identifying selected loci in E&R studies (Iranmehr et al., 2017). We used allele frequency estimates of each $10^th$ generation (10 time points in total with 90 generations of selection). Since CLEAR is computationally demanding we analyzed the data solely for one chromosome arm. The selection regime had a significant influence on the power to identify QTNs (Kruskal–Wallis rank-sum test with pAUC=0.01; $p = 5.79e - 05$). Furthermore, the increasing regime had a better performance than the constant regime (Wilcoxon rank sum test with pAUC=0.002; $90 \rightarrow 10\%$ vs 50%; $p = 0.035$).

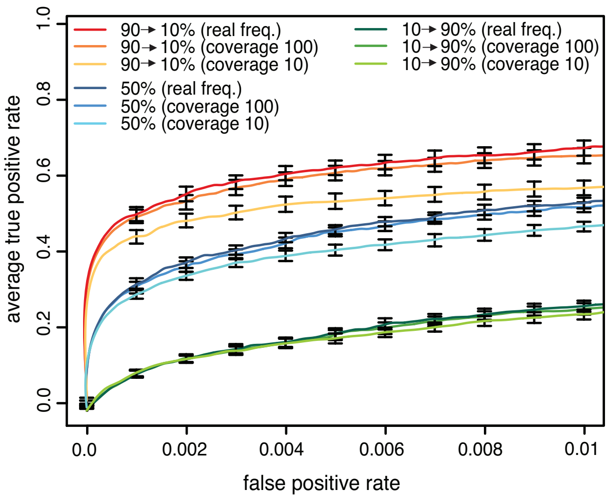


Figure 18: Influence of the coverage of Pool-Seq data on the power to identify QTNs. For each selection regime we show the performance using the actual allele frequency and allele frequency estimates obtained with Pool-Seq . Binomial sampling was used to model Pool-Seq with coverages 100 and 10. As expected the performance of E&R studies decreases with the coverage (Kofler and Schlötterer, 2014). Nevertheless the increasing regime outperformed the constant regime irrespective of the coverage (Wilcoxon rank sum test with pAUC; $90 \rightarrow 10\%_{realfreq}$ vs $50\%_{realfreq}$; $p = 2.165e{-}05$; $90 \rightarrow 10\%_{cov.100}$ vs $50\%_{cov.100}$; $p = 2.165e{-}05$; $90 \rightarrow 10\%_{cov.10}$ vs $50\%_{cov.10}$; $p = 2.165e - 05$).

Table 1: Overview of the three best selection regimes for different experimental designs and trait architectures. Selection regimes were ranked based on the partial area under the ROC curve ($pAUC$). Starting from the default conditions (top) we varied one parameter at the time (bold). g number of generations, r replicates, n number of loci, $h^2$ heritability, N population size, ga. gamma distributed effect sizes , eq. equal effect sizes

| parameters | best regimes | pAUC±sd |
|---|---|---|
| g=90,r=10,n=100 $h^2$=1,N=1000,ga. | 90 → 20% | 0.00638 ± 0.0005 |
| | 90 → 30% | 0.00633 ± 0.0004 |
| | 90 → 40% | 0.00622 ± 0.0004 |
| **g=20**,r=10,n=100,$h^2$=1,N=1000,ga. | 90 → 70% | 0.00362 ± 0.0004 |
| | 90 → 60% | 0.00356 ± 0.0003 |
| | 80% | 0.00352 ± 0.0003 |
| g=90,**r=5**,n=100,$h^2$=1,N=1000,ga. | 90 → 30% | 0.00540 ± 0.0004 |
| | 90 → 40% | 0.00529 ± 0.0004 |
| | 90 → 20% | 0.00524 ± 0.0003 |
| g=90,r=10,**n=1000**,$h^2$=1,N=1000,ga. | 90 → 20% | 0.00157 ± 0.0001 |
| | 90 → 10% | 0.00154 ± 0.0001 |
| | 90 → 30% | 0.00153 ± 0.0001 |
| g=90,r=10,n=100,**$h^2$=0.6**,N=1000,ga. | 90 → 10% | 0.00450 ± 0.0003 |
| | 90 → 40% | 0.00449 ± 0.0003 |
| | 90 → 20% | 0.00448 ± 0.0004 |
| g=90,r=10,n=100,$h^2$=1,**N=2000**,ga. | 90 → 30% | 0.00682± 0.0004 |
| | 90 → 10% | 0.00670 ± 0.0005 |
| | 90 → 20% | 0.00669 ± 0.0004 |
| g=90,r=10,n=100,$h^2$=1,N=1000,**eq.** | 90% | 0.00915 ± 0.0002 |
| | 90 → 80% | 0.00828 ± 0.0005 |
| | 90 → 70% | 0.00759 ± 0.0002 |

Table 2: Performance of the 90 → 20% selection relative to the regime with the best performance with a given set of parameters. We used pAUC to assess the performance of a regime and compute the ratio between the 90 → 20% increasing regime and the best regime (br). A value of 74%, for example, indicates that the 90 → 20% increasing regime is 26% less powerful than the best regime. g number of generations, r replicates, n number of loci, $h^2$ heritability, p population size, ga. gamma distributed effect sizes with shape 0.42, ga.1 gamma distributed effect sizes with shape 0.1, ga.2 gamma distributed effect sizes with shape 0.7, ga.3 gamma distributed effect sizes with shape 1.0, eq. equal effect sizes

| parameters | best regime | $pAUC_{90\rightarrow 20}/pAUC_{br}$ (%) |
|---|---|---|
| **g=90**,r=10,n=100 $h^2$=1,p=1000,ga. | 90 → 20% | 100 |
| **g=45**,r=10,n=100,$h^2$=1,p=1000,ga. | 80 → 10% | 99 |
| **g=20**,r=10,n=100,$h^2$=1,p=1000,ga. | 90 → 70% | 85 |
| g=90,**r=5**,n=100,$h^2$=1,p=1000,ga. | 90 → 30% | 97 |
| g=90,**r=3**,n=100,$h^2$=1,p=1000,ga. | 80 → 50% | 97 |
| g=90,r=10,**n=1000**,$h^2$=1,p=1000,ga. | 90 → 20% | 100 |
| g=90,r=10,**n=25**,$h^2$=1,p=1000,ga. | 90 → 20% | 94 |
| g=90,r=10,n=100,**$h^2$=0.6**,p=1000,ga. | 90 → 10% | 99 |
| g=90,r=10,n=100,**$h^2$=0.3**,p=1000,ga. | 90 → 20% | 100 |
| g=90,r=10,n=100,$h^2$=1,**p=2000**,ga. | 90 → 30% | 98 |
| g=90,r=10,n=100,$h^2$=1,p=1000,**ga.1** | 90 → 20% | 100 |
| g=90,r=10,n=100,$h^2$=1,p=1000,**ga.2** | 90 → 30% | 97 |
| g=90,r=10,n=100,$h^2$=1,p=1000,**ga.3** | 90 → 40% | 97 |
| g=90,r=10,n=100,$h^2$=1,p=1000,**eq.** | 90% | 74 |

### References

H. Bastide, A. Betancourt, V. Nolte, R. Tobler, P. Stöbe, A. Futschik, and C. Schlötterer. A Genome-Wide, Fine-Scale Map of Natural Pigmentation Variation in *Drosophila melanogaster*. *PLoS Genetics*, 9: e1003534, 2013.

J. Comeron, R. Ratnappan, and S. Bailin. The Many Landscapes of Recombination in *Drosophila melanogaster*. *PLoS Genetics*, 8(10):e1002905, 2012.

A. Iranmehr, A. Akbari, C. Schlötterer, and V. Bafna. CLEAR: Composition of likelihoods for evolve and resequence experiments. *Genetics*, 206(2):1011–1023, 2017.

R. Kofler and C. Schlötterer. A Guide for the Design of Evolve and Resequencing Studies. *Molecular biology and evolution*, 31(2):474–483, 2014.

K. Spitzer, M. Pelizzola, and A. Futschik. Modifying the Chi-square and the CMH test for population genetic inference: adapting to over-dispersion. 2019. URL http://arxiv.org/abs/1902.08127.
